# Reaching the full potential of cryo-EM reconstructions with molecular dynamics simulations at 310 K: Actin filaments as an example

**DOI:** 10.1101/2025.08.11.669737

**Authors:** Sahithya Sridharan Iyer, Kristina M. Herman, Yihang Wang, Thomas D. Pollard, Gregory A. Voth

## Abstract

Cryo-electron microscopy (cryo-EM) structures of multi-protein complexes such as actin filaments help explain the mechanisms of assembly and interactions with partner proteins. Yet, rapid cooling during freezing may not preserve the conformations at physiological temperature. All-atom molecular dynamics simulations starting with cryo-EM reconstructions can provide additional insights. For example, at 310 K the states of ADP-actin filaments consistent with higher entropy favor partly twisted subunits and smaller rotations along short-pitch helix than the cryo-EM reconstructions, while cryogenic temperatures favor flattened conformations. In the active site, the positions of Q137 and the catalytic water 1 and activating water 2 optimal for in line attack on the γ-phosphate of ATP are very rare at 310 K, explaining in part the slow rate of ATP hydrolysis in filaments. This favorable arrangement of the waters is not observed in simulations of actin monomers. At 310 K subunits in ADP-P_i_-actin filaments have their backdoor gates open 60% of the time for phosphate release, a conformation not observed by cryo-EM. Rare fluctuations open binding sites for cofilin and phalloidin. The twisted conformations of pointed end subunits and interactions of the D-loop of the penultimate subunit explain the slow association of new subunits. The terminal subunit at the barbed end is tethered to its neighbor along the long-pitch helix but transiently dissociates from its lateral neighbor. These effects of subfreezing temperatures on actin filaments are surely not an isolated example, so MD simulations of structures of other frozen proteins will be informative.

**Significance statement:** Functionally important properties of proteins are tightly linked to their conformations at physiological temperatures. While cryo-EM reconstructions and crystal structure of frozen proteins provide a molecular resolution of protein structures, they might differ from the conformations at physiological temperature. Using the actin filament as a case study, we find cryo-EM reconstructed structures correspond to low entropy conformations that differ from the ensemble of structures in molecular dynamics simulations at 310 K. The fluctuations of subunit dihedral angles, short-pitch rotations and some side chains explain functionally important properties of actin filaments, including the slow rates of ATP hydrolysis and phosphate release as well as the slow binding of the protein cofilin and cyclic peptide phalloidin.

## Introduction

Understanding the structure and dynamics of biomolecules in their native environment has been an enduring challenge of molecular biophysics. Among the structure determination methods, cryo-electron microscopy (cryo-EM) stands out as it circumvents the need to grow single crystals while resolving structures at atomic resolution (Bai, McMullan and Scheres 2015, Nakane, Kotecha et al. 2020, Nogales and Mahamid 2024).

Actin filaments are not only the main component of the cytoskeleton but also essential for cell movements. Monomers of actin self-assemble head to tail into filaments (Pollard and Cooper 2009) with two strands of subunits in a double-helix, with all the subunits oriented in the same direction and stabilized by inter-stand (lateral) and intra-stand (longitudinal) interactions (Fig. 1A). Structural differences at the two ends of actin filaments result in kinetic polarity in polymerization, with the barbed (plus) end polymerizing faster than the pointed (minus) end (Pollard 1986, Zsolnay, Katkar et al. 2020).

**Figure 1:**
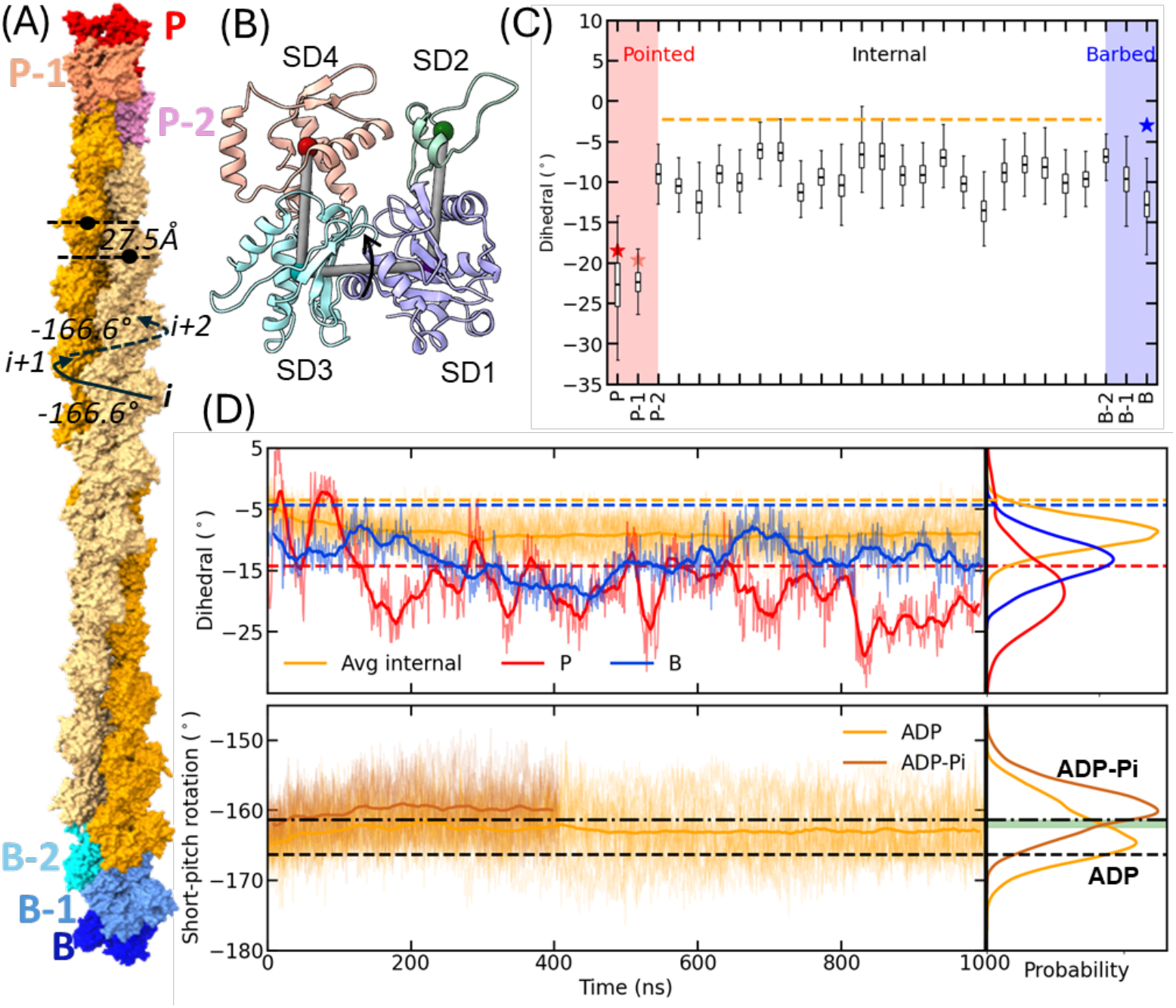
Actin subunits at the ends and middle of the filament become more twisted during MD simulations of a filament with 27 ADP-actin subunits at physiological temperature of 310 K. (A) Actin filament constructed by stitching together cryo-EM structures of a free pointed end (pdb 8F8S), internal subunits (pdb 8F8P) and free barbed end (pdb 8F8R). The colored labels name subunits. The black labels and curved arrows define the left-handed short-pitch helix with a rotation angle of -166.6° and a rise of 27.5 Å. (B) Ribbon diagram the actin monomer with four subdomains: SD1 (purple), SD2 (green), SD3 (cyan) and SD4 (pink). Centers-of-mass of Cα atoms of the subdomains were used to calculate the dihedral angle SD2-SD1-SD3-SD4. (C) Dihedral angles of each subunit calculated during the last 200 ns of the 1 μs simulation. Edges of the box plots represent first and third quartiles and the horizontal lines within each box are the mean dihedral value. The whiskers extend to minimum and maximum dihedrals sampled in the last 200 ns of the simulation. The horizontal dashed line and the stars are the dihedral angles in the starting structure. (D) Time courses of (top) dihedral angles and (bottom) short-pitch helical rotations during the simulations with probability distributions to the right. Darker lines are running averages calculated in 20 ns windows. Lighter shaded lines are raw data. (Top) Barbed and pointed end subunits and average of internal subunits. Horizontal dashed lines are the values in the cryo-EM reconstructions. (Bottom) Short-pitch rotation angles for ADP-(orange) and ADP-P_i_-(brown) actin filaments. Horizontal black lines are Short-pitch rotation angles in cryo-EM reconstructions of (dashed) an ADP-actin filament (pdb id: 8F8P) and (dot dashed) cofilin bound ADP-actin filament (pdb id: 5YU8). Probability distributions of helical rotation angles in the last 500 ns of the ADP-actin filament and 200 ns of the ADP-P_i_-actin filament. The green bar indicates angles of - 161.4° to -162.4° that are favorable for binding cofilin.

The actin molecule has an outer domain comprised of subdomains SD1 and SD2 and a larger inner domain composed of SD3 and SD4 (Kabsch, Mannherz et al. 1990) (Fig. 1B). X-ray fiber diffraction of well-oriented filaments at 4°C revealed (Oda, Iwasa et al. 2009) and cryo-EM (Fujii, Iwane et al. 2010, Merino, Pospich et al. 2018, Chou and Pollard 2019, Carman, Barrie et al. 2023) confirmed that the subunits are flattened by rotation of the outer domain about 15° with respect to the inner domain compared with actin monomers . This conformational change is measured by the dihedral angle formed by SD2-SD1-SD3-SD4. Along the one-start, short-pitch helix including all the subunits, each successive subunit is rotated -166° relative to its neighbor. Flattening actin subunits during polymerization and reorientation of side chains in the active site increases the rate of ATP hydrolysis 42,000-fold (Blanchoin and Pollard 2002, Rould, Wan et al. 2006, McCullagh, Saunders and Voth 2014, Sun, Sode et al. 2017).

Cryo-EM reconstructions of actin filaments confirmed that the internal and barbed end subunits are flattened, while the pointed end subunits are more twisted similar to actin monomers (Carman, Barrie et al. 2023). Narita et al. (Narita, Oda and Maéda 2011), Zsolnay et al. (Zsolnay, Katkar et al. 2020) and Carman et al. (Carman, Barrie et al. 2023) proposed the twisted pointed end subunits with the subunit P-1 D-loop interacting with subunit P make the end unfavourable for the addition of an actin subunit, explaining slow elongation at the pointed end.

In this study and in the most general sense, we focused on how physiological temperature may cause multi-protein complexes to deviate from their structures determined at cryogenic temperature after rapid freezing. Prior pioneering work explored this issue from a different perspective with the ribosome ⋅ EF-Tu complex as the example (Bock and Grubmüller 2022). Taking actin filaments as our example, we used all-atom molecular dynamics (MD) simulations at 310 K starting with cryo-EM reconstructions of ATP-, ADP-P_i_- and ADP-actin filaments. The subunits in simulated actin filaments with 7, 13 or 27 subunits fluctuated around more twisted average conformations with smaller helical rotations than the starting cryo-EM structures. The flattened cryo-EM structures (but analyzed at 310 K) have lower entropies and higher free energies than the MD structures at that temperature, with the MD structure therefore satisfying the well-established Principle of Maximum Entropy (Jaynes 1957, Jaynes 1957). Fluctuations of sidechain rotameric conformations and dynamics of coordinated water molecules at 310 K account for the slow the rates of ATP hydrolysis and phosphate release. Mutation of backdoor gate residue N111 to S increases the rate of P_i_-release due to the reduced probability of closed backdoor gates and “occluded states” of the phosphate release channel, which, in turn, influences the positioning of H161 in the active site (Wang, Wu et al. 2024). Rare fluctuations of the filament are required to bind cofilin and phalloidin. The simulations confirm that pointed end subunits are more highly twisted with the D-loop of subunit P-1 interacting with subunit P. We also confirm that the terminal subunit at the barbed end is twisted and anchored to subunit B-2 by its D-loop but transiently loses other interactions with subunit B-1 and B-2 (see Fig. 1A and next section for definitions).

## Results

### Cryo-EM reconstructions used to model a filament with 27 ADP-actin subunits

We joined cryo-EM reconstructions (Carman, Barrie et al. 2023) of a free pointed end (pdb id: 8F8S), internal subunits (pdb id: 8F8P) and a free barbed end (pdb id: 8F8R) to make an ADP-actin filament with 27 subunits to start the all-atom MD simulations at 310 K (Fig. 1A and Movie S1). To describe the conformations of the subunits we use the dihedral angles SD2-SD1-SD3-SD4 (Fig. 1B) of the subunits and the rotation between subunits along the short-pitch helix (Fig. 1A). As explained in the original structure determinations (Carman, Barrie et al. 2023), the internal subunits (I) in this model have dihedral angles -2.3° and a short-pitch rotation angle of -166.6°, and the terminal subunit at the barbed end (B) has a dihedral angle of -3.0° and a short-pitch rotation angle of -166.1°, both flattened conformations compared with the twisted conformation of the actin monomer (−20.3°). At the pointed end of the model, the terminal subunit P has a dihedral angle of -18.5° and the penultimate subunit P-1 has a dihedral angle of -19.6° and a short-pitch rotation angle of -166.3°, both nearly as twisted as an actin monomer.

### All-atom MD simulations of the 27-mer ADP-actin filament at 310 K

As expected, the conformations of all the subunits in actin filaments fluctuated during 1 μs simulations at 310 K (Supporting Information Movies S1, S2 and S3), but unexpectedly, the conformations fluctuated around different mean values (Fig. 1C, D) than the static conformation captured in the starting cryo-EM reconstructions (horizontal dashed lines in Fig. 1C, D). The transitions to more twisted conformations with larger short-pitch helical rotations occurred during the first 200-300 ns of simulation (Fig. 1D, yellow curves). Two additional replica simulations of 450 ns gave similar results (Fig. S1). Tables S1 and S2 compare the subunit dihedral angles and helical rotations in the cryo-EM reconstructions and MD simulations.

Movies S1, S2 and S3 with images every 10 ns show the main features of the “warm” ADP-actin filament at 310 K: (a) interactions between the D-loop of subunit *i* with subdomain 3 of subunit *i+2* form stable longitudinal interactions; (b) lateral interactions between internal subunits open up transiently, and the lateral inactions of subunit B with subunit B-1 dissociate and reform (Movies S2 and S3); (c) the D-loop of subunit P (red) fluctuates freely in the solvent; (d) the D-loop of subunit P-1 (tan) explores the surface of subunit P; (e) fluctuations in the helical rotations between subunits along the short-pitch helix cause the filament to twist and bend locally; and (f) the filament is stiff with a persistence length of 8.2 μm in the 1 μs simulation (Fig. S2), as experimentally observed with filaments in solution (Ott, Magnasco et al. 1993, Isambert, Venier et al. 1995).

The next section describes why the filaments change during MD simulations at 310 K owing to the effects of temperature on the conformations, free energies and conformational entropies of twisted and flattened actin subunits at 310 K. The following sections also document how the fluctuations of conformations of internal subunits influence ATP hydrolysis, phosphate dissociation and binding of cofilin and phalloidin. The final sections describe the behavior of filament ends at 310 K.

### Temperature dependence of the conformations of actin filaments during MD simulations

We simulated ADP-actin filaments at 90 K, 175 K, 230 K, 250 K, 275 K and 298 K for comparison with simulations at 310 K (Figs. 2B, S3 and S4). As expected, the dihedral angles and helical rotations of all 27 subunits in the model filament were constant during an MD simulation at the cryo-EM temperature of 90 K (Table S1). A simulation of a 13-mer ADP-actin filament at 90 K was the same. Note that the dihedral angles and helical rotations changed modestly from the initial cryo-EM model during the initial energy minimization process (Tables S1 and S2) but deviated minimally thereafter during the production simulation at 90K as indicated by the small standard deviations in the angles.

**Figure 2:**
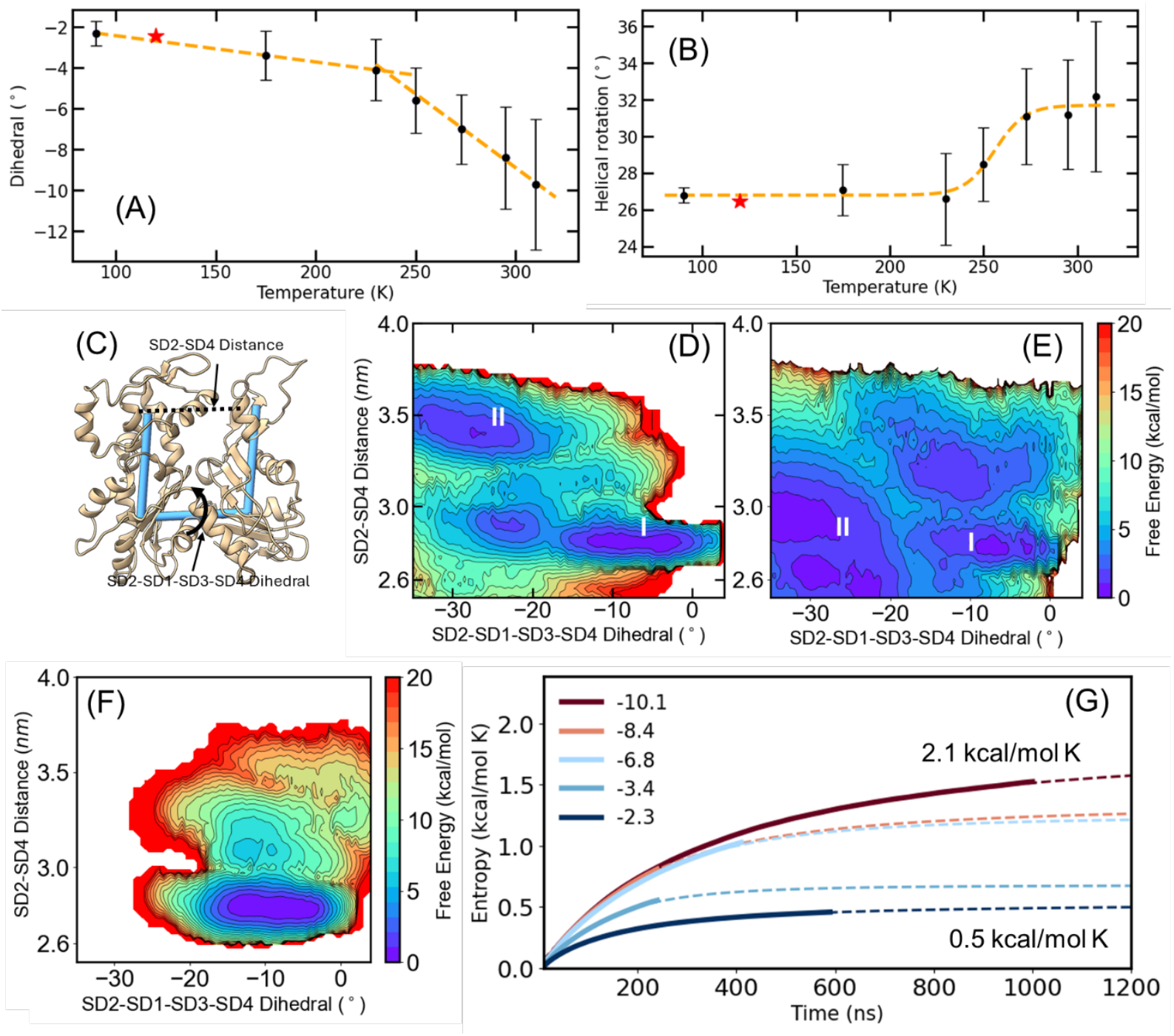
Thermodynamics of conformational changes in actin filament subunits. Temperature dependence of average (A) subunit dihedral angles and (B) helical rotation angles of internal subunits in an ADP-actin filament with 13 subunits. Error bars are standard deviations. Dashed lines are eye guides. Stars indicate the dihedral angle and helical rotation in cryo-EM reconstruction (pdb id: 8D18) refined by MD simulation at 120 K. (C) Ribbon diagram of an actin subunit showing two collective variables (CVs) used in the well-tempered metadynamics simulation of a seven-subunit actin filament. CV1 is the dihedral angle formed by SD2-SD21-SD3-SD4 and CV2 is the distance between SD2 and SD4. (D-F) Two dimensional free energy landscape surfaces as a function of dihedral angle and distance between SD2 and SD4 at 310 K. (D) Barbed end subunit, (E) pointed end subunit and (F) internal subunits. (G) Time courses of calculations of backbone conformational entropy using the Schlitter approximation of backbone atoms of internal actin subunits of an ADP-actin filament with 13 subunits with 5 different average dihedral angles shown in the inset. The dashed lines correspond to the fit to a sigmoid-like function to approximate entropy values over longer times.

Simulations of the 13-mer ADP-actin filament at temperatures between 90 K and 310 K showed that the internal subunits were more twisted (more negative dihedral angles) with larger helical rotation angles than the starting cryo-EM model (Fig. 2A,B). The standard deviation bars document that the fluctuations in the dihedral and helical rotation angles increased with temperature as expected. Below a threshold temperature of ∼250 K, the subunits were less twisted like cryo-EM reconstructions (Fig. 2A). The small slope of the interpolated line between temperatures 90 K and 230 K reflects the small changes in the structure in this temperature range. Molecular Dynamics Flexible Fitting (MDFF) (Trabuco, Villa et al. 2008) refinement at an upper temperature of 120 K of a cryo-EM reconstruction of actin filaments (pdb id: 8D18 (Reynolds, Hachicho et al. 2022)) produced subunit dihedral and helical rotation angles matching our interpolated lines (Fig. 2 A,B red stars).

### Free energies of twisted and flattened actin subunits at 310 K

We used well-tempered metadynamics (WTMetaD) (Barducci, Bussi and Parrinello 2008, Dama, Parrinello and Voth 2014) simulations to calculate the free energies at 310 K associated with twisting of subunit P, subunit B and internal actin subunits in filaments with seven ADP-actin subunits. We used two collective variables (CVs) to define these motions: the dihedral angle and the distance between SD2 and SD4 (Fig. 2C).

Internal subunits with dihedral angles in the range -17° to -3° were conformations with similar free energy minima (Fig. 2F). Conformations outside this range, with dihedral angles less than -18° and greater than -2°, had unfavorable free energies.

The barbed end terminal subunit B had two broad free energy minima (blue areas I and II, Fig 2D). Energy basin I has two regions with the B subunit tethered to B-2 by the D-loop and β-turn residues P243-Q246 in SD4. The first minimum between dihedral angles of -20° to -3°. A second minima between -20 and -28°, to the left of I, connected to I by barriers less than 3 kcal/mol, corresponds to conformations of the B subunit with a twisted dihedral angle but remaining in contact with subunit B-2 though SD3 contacts. In free energy basin II subunit B was extremely twisted with dihedral angles between -30° to -20°, while being tethered to subunit B-2 by the D-loop without longitudinal contacts with B-2 via the β-turn mediating the SD3/SD4 contacts or lateral contacts with B-1. Pointed end terminal subunit P had a broad free energy basin with twisted (I) and flat (II) conformations (Fig. 2E) separated by a barrier of less than 1 kcal/mol. Consequently, they easily interconvert.

### Entropic preference for twisted over flattened actin subunits at 310 K

To investigate why actin subunits in cryo-EM structures adopt flatter conformations than in simulations at higher temperatures despite having similar free energies, we used the Schlitter quasi-harmonic approximation to calculate the conformational entropy of backbone atoms of actin in an ADP-filament with 13 subunits over a range of dihedral angles (Fig. 2G). The backbone conformational entropies were calculated from the covariance matrix of atomic fluctuations (Methods, Eqn 1) (Schlitter 1993, Schäfer, Mark and van Gunsteren 2000).

The methods section explains that the Schlitter entropy is an upper bound to the true conformational entropy as it does not account for the anharmonicity and higher order correlations in fluctuations. The Schlitter approximation, rather than absolute values. is permissible for our calculations of the difference in the entropies of twisted and flat conformations at 310 K. We removed rigid body motion corresponding to translation and rotation by calculating the covariance matrix for trajectories that are aligned to the reference configuration. The entropy of internal subunits with different average dihedral angles is calculated by scaling the fluctuations measured at the respective temperatures to 310 K (Fig 2G). The build ups in entropy during the simulations were fit to a sigmoid-like function, 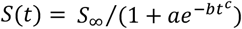where *S*_∞_, *a, b* and *c* are fit parameters, to estimate the asymptotic time entropy values.

These calculations revealed that the entropy of the most twisted subunits with an average dihedral angle of -10.1° is about 4 times larger than the flattest subunits with an average dihedral angle of -2.3° (Fig. 2G). By drawing upon the well-established Principle of Maximum Entropy (Jaynes 1957, Jaynes 1957), i.e., that at a given temperature a system at equilibrium will always achieve the maximum entropy state, this difference in entropy explains why higher entropy twisted conformations are preferred at 310 K and lower entropy flattened conformations become favored as a sample of filaments approach 273 K during freezing of cryo-EM samples. At temperatures below 273 K, the flat subunit conformation has a lower free energy that becomes more enthalpically favored with lower entropy, whereas at higher temperatures above 273 K, the twisted conformation has a lower free energy that becomes more entropically stabilized (Fig. 2G). Simply put, the twisted state at 310 K explores more conformational fluctuations and hence has a higher entropy than the flattened state, which is more constrained.

### Conformations of internal subunits in actin filaments during MD simulations at 310 K

During the all-atom MD simulations at 310 K, every subunit in the ADP-actin filament transitioned to more twisted conformations with larger (less negative) rotations of the short-pitch helix than the starting cryo-EM reconstructions (Fig. 1C, D). After 200-300 ns of simulation the average dihedral angles of the 23 internal subunits plateaued at -9° ±2° (± SD) (Fig. 1D, yellow curves) as they sampled a wide range of dihedral angles shown by the whiskers in Fig. 1C and fine lines around the mean angles (Fig. 1D, top).

The short-pitch helical rotations between internal subunits i and i+1 changed from -166.6° in the cryo-EM model (Carman, Barrie et al. 2023) to an average of -163° during the 1 μs simulation, while sampling a large range of angles between -176° to -150° (Fig. 1D, bottom). These rotations were largely uncorrelated with the dihedral angles (Fig. S5A). Thermal fluctuations cause small bends over time and allow transient losses of lateral interactions between internal subunits (Movie S1) as noted by electron microscopy of negatively stained filaments (Bremer, Millonig et al. 1991); however, the loss of lateral interactions was only weakly correlated with the dihedral angles and helix rotation angles (SI Section 4, Fig. S5).

The short-pitch rotations in ADP-P_i_-actin filaments changed from -166.5° in the cryo-EM model (pdb id: 8A2S) (Oosterheert, Klink et al. 2022) to -160° during the 380 ns simulation at 310 K, while sampling a large range of angles between -170° to -148° (Fig. 1D, bottom). Cryo-EM reconstructed structures of ADP- and ADP-P_i_-actin filaments did not have this 3° difference in short-pitch rotations (Chou and Pollard 2019, Oosterheert, Klink et al. 2022).

### Effects of subunit dynamics at 310 K on ATP hydrolysis

High resolution crystal (pdb id. 7W4Z; (Kanematsu, Narita et al. 2022, Iwasa, Takeda et al. 2023)) and cryo-EM (pdb id: 8A2R; (8)) structures established the requirements (Fig. 3A) for inline attack of catalytic water 1 to hydrolyze the γ phosphate from ATP (McCullagh, Saunders and Voth 2014, Sun, Sode et al. 2017, Oosterheert, Klink et al. 2022). First, Q137 must position catalytic water 1 ∼ 4 Å from the phosphate. Second, water-1 must be nearly in line with the phosphate with an O-γ phosphate-W1 angle >150° (Scipion, Ghoshdastider et al. 2018). Third, the oxygen of W1 must be oriented toward the phosphate. Fourth, a second water, water 2 (W2), must be positioned close to W1 to allow for hydrogen bonding and activation of W1.

**Figure 3:**
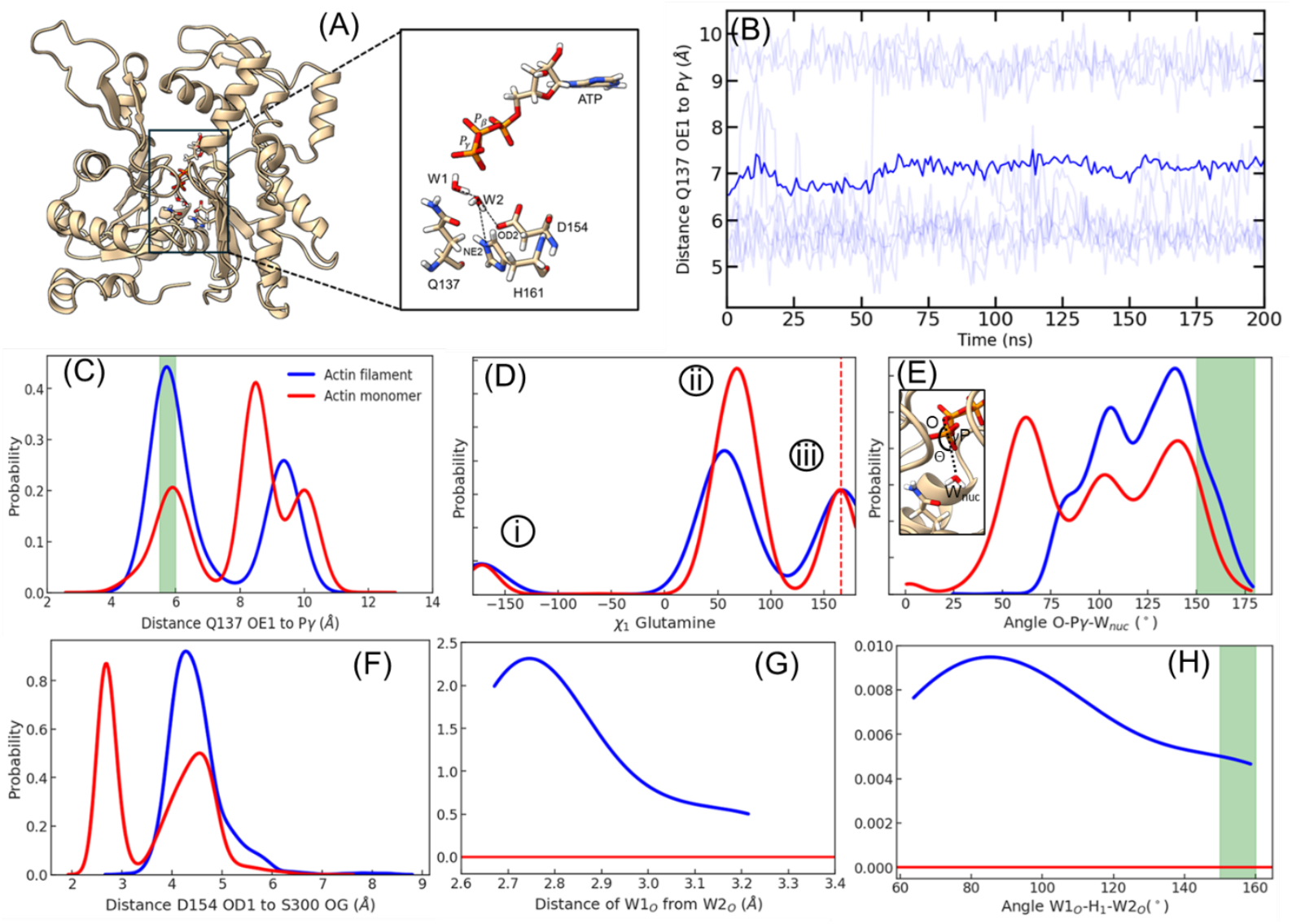
Motions in the active site required for ATP hydrolysis during 200 ns simulations at 310 K of an ATP-actin filament with 13 subunits and 4 μs for an ATP-actin monomer. Nucleophilic attach of water 1 to hydrolyze ATP depends on the positions of residues Q137, H161 and D154 and waters 1 (W1) and 2 (W2). (A) Ribbon diagram of an internal actin filament subunit from the simulation with Q137, D154, H161, ATP, W1 and W2 in CPK representation with dotted lines for hydrogen bonds. (B) Time evolution of the distance between Q137 OE1 and the γ phosphate of ATP in internal subunits of a 13-mer ATP-actin filament. Light traces are distances in individual internal subunits and the dark trace is the average across all the internal subunits. (C-H) Probability distributions of interatomic distances, side chain dihedral and bond angles in internal subunits of an ATP-actin filament (blue) and ATP-actin monomer (red). Green shaded regions mark distances and angles suitable for catalysis. (C) Distances between Q137 OE1 and γ phosphate. (D) Probability distribution of *χ*_1_ rotamers of Q137. Dashed line indicates the *χ*_1_ angle in cryo-EM reconstruction pdb id 8A2R. (E) The angle O-γ phosphate-W1 for subunits with distances of 5.5 Å to 6.0 Å between Q137 OE1 and γ phosphate. (F) Probability distribution of distance between OD1 of D157 and OD of S300. (G) Probability distribution of distance between W1 and W2. W2 is identified as a water located within 3.5 Å of W1 oxygen and coordinated to D157 and H161. No W2 were identified for monomer. (H) Probability distribution of the angle OW1-HW1-OW2. Angles that are optimal for hydrogen bonding are marked in green shaded region.

The arrangement of Q137 and the two crucial waters in the active site changed with fluctuations of subunit dihedral angles (Figs. 3B-H and S11) (McCullagh, Saunders and Voth 2014, Sun, Sode et al. 2017, Oosterheert, Klink et al. 2022). For example, distance between OE1 of Q137 to γ phosphate of ATP sampled a bimodal distribution (Fig. 3B, C) with peak positions at the catalytic distance of ∼5.5-6 Å and at non-catalytic distance of 9.5-10 Å. No waters were observed at distances less than 5 Å. The variation between catalytic and non-catalytic distances correspond to 3 *χ*_1_ rotameric states of Q137 at 310 K (Fig. 3D, S6).

We employed MD simulations at 310 K to measure the fraction of time when these conditions for hydrolysis were met in Mg-ATP-actin filaments with 13 subunits and ATP-actin monomers. Each of the four requirements for hydrolysis impacted the overall reaction. Condition 1: OE1 of Q137 was positioned 5.5 to 6.0 Å from γ phosphate of ATP suitable to position the catalytic W1 ∼ 4 Å form the phosphate during a fraction of 0.32 of the time in interior subunits of filaments and 0.11 of the time in actin monomers (Fig. 3C). Both rotameric states (iii) with χ1 of 160° and (i), with χ1 of -165°.are suitable for catalysis when positioned between 5.5 to 6.0 Å from γ phosphate of ATP (Figs. 3D and S6,S7). Condition 2: Of the conformations that satisfied condition 1, only 0.15 of interior subunits in filaments and 0.07 of actin monomers had O-γ phosphate-W1 angles >150° as required for efficient nucleophilic attack (Fig. 3E). Condition 3: If either HW1 or HW2 of W1 forms an angle less than 90° with the vector pointing from water 1 oxygen (OW) to PG, the hydrogen points toward PG, so the orientation of W1 is unfavourable for catalysis. Of the conformations that satisfied conditions 1 and 2, 0.5 of waters in interior subunits and 0.3 of waters in monomeric actin had the oxygen of the attacking W1 with orientation suitable for catalysing ATP hydrolysis. Condition 4: W2 must be positioned within H-bonding distance of W1 to facilitate the deprotonation of W1 during the nucleophilic attack (Fig. 3F-H). W2 is within 3.5 Å of W1, OD2 of D156 and ND2 of H161 during a fraction of 0.02 of the time in interior subunits of filaments. During this time, W2 has an optimal geometry to hydrogen bond with W1 for a fraction of 0.25 of the time in actin filament subunits (Fig. 3 G,H) .

Combining these probabilities, the fraction of subunits in Mg-ATP-actin filaments with catalytically favorable waters is 0.32 x 0.15 x 0.5 x 0.02 x 0.25 = 0.00012. These rare subunits hydrolyze ATP at 0.3 s^-1^/0.00012 = 2500 s^-1^.

The 4 μs simulation of a Mg-ATP-actin monomer never sampled the full set of conditions in the active site required for hydrolysis. The first three conditions occurred frequently with probabilities of 0.11, 0.07 and 0.3 (Figs. 3C,E), but W2 in the correct position was not observed (Fig. 3G,H), explaining the very slow hydrolysis rate (Rould, Wan et al. 2006). Either D154 was hydrogen bonded to S300 precluding placement of W2 (Fig. 3F) and/or D154 separated from S300 hydrogen bonded to W2 at an angle inappropriate to hydrogen bond W1.

### Effects of subunit dynamics at 310 K on phosphate release pathway

In most, but not all (Chou and Pollard 2023), cryo-EM structures of ADP-P_i_ and ADP-actin filaments (for example, pdb id: 8A2S and 8A2T) (Oosterheert, Klink et al. 2022) a hydrogen bond between the sidechains of N111 and R177 forms a “backdoor gate” that helps to block phosphate release from ADP-P_i_-actin filaments (Oosterheert, Blanc et al. , Wang, Wu et al. 2024, Oosterheert, Boiero Sanders et al. 2025)., but this gate opened reversibly for a majority of the time during our MD simulations (Fig. 4A,B). During the simulations, the probabilities of open gates with separations >5 Å was 0.7 for filaments with 27 ADP-actin subunits and 0.6 for filaments with 27 ADP-P_i_-actin subunits.

**Figure 4:**
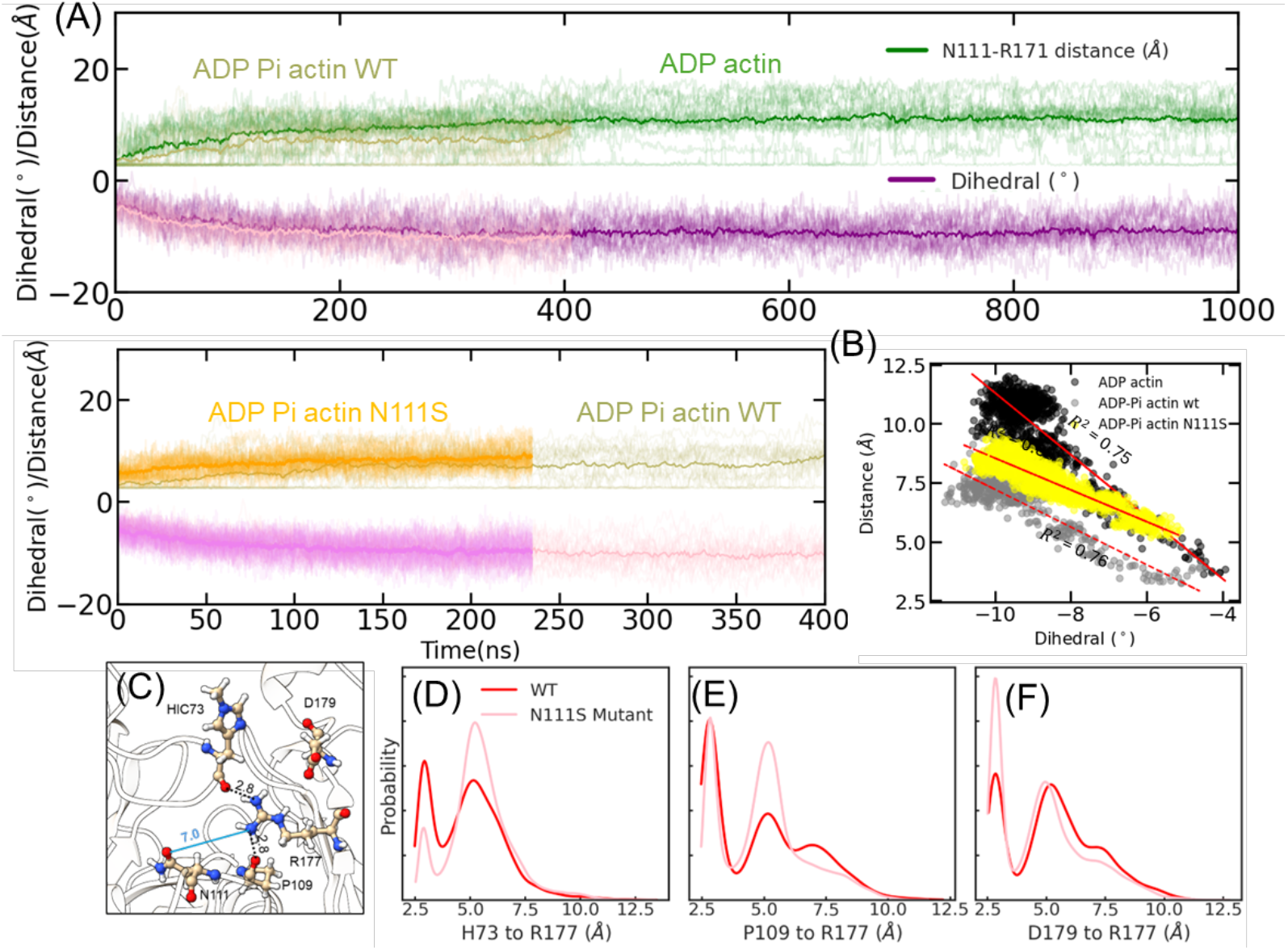
Dynamics of the phosphate release pathway during simulations. (A) Time evolution of dihedral angles and the minimum distances between OD1 of S111/OG of N111 and NH1 and NH2 of R177 of all 23 internal subunits of the (Top) ADP-(darker shade) and ADP-P_i_-filaments (lighter shade) and (Bottom) ADP-P_i_-WT (khaki/light pink) and ADP-P_i_-N111S mutant (orange/pink) actin filaments. The darker lines are averages across the internal subunits. Lighter traces are dihedral angles and distances of individual internal subunits. Only 2 of the 23 internal subunits have a closed backdoor (separation <4 Å) for more than 40% of the 1 μs simulation. (B) Plot of the average distance measured between N111/S and R177 in (A) vs. average dihedral angle of the subunits during the simulation of an ADP, ADP-Pi and ADP-Pi-N111S actin filament. (C) CPK representation of residues H73, P109, N111, R177 and D179 in the occluded state. Dashed black lines are hydrogen bonds; blue lines are non-bonding distances. (D-F) Probability distributions of minimum distances between NH1/NH2 of R177 and interacting atoms in the wild type and mutant actins: (D) backbone oxygen of H73; (E) backbone oxygen of P109; and (F) OD1/OD2 of D179.

The distance between the side chains forming the gate was tightly coupled to the fluctuations of the dihedral angles formed by SD2-SD1-SD3-SD4 (Fig. 4B), because N111 in SD1 and R177 in SD3 move relative to each other as the dihedral angle changes. The absence of the γ-phosphate in the active site of ADP-actin filaments favored the formation of a salt bridge between R177 and D179, which held the backdoor gate open. This, in turn, allowed wider separation of N111 and R177, explaining why the slope in Fig. 4B is steeper for ADP-actin than ADP-P_i_-actin filaments. The hydrogen bond between N111 and R177 is not strong enough to resist thermal motions, so subunit twisting reversibly separates these residues to open and close the backdoor gate.

Sidechain fluctuations in the ADP-P_i_-actin filament generated a rapidly fluctuating network of hydrogen bonds between the sidechain of R177 and the backbone carbonyl oxygens of H73, P109 and H161 in the phosphate exit channel that blocks diffusion of phosphate (Fig. 4 C-F). In the occluded state N111 is hydrogen bonded to E107. This “occluded state” was discovered in prior MD simulations (Wang, Wu et al. 2024) but never seen in cryo-EM structures.

Mutation of N111 to serine increases the rate of phosphate release about 15-fold (Oosterheert, Blanc et al. 2023), so we simulated N111S ADP-P_i_ actin filaments with 27 subunits to investigate the mechanism. At steady state the fraction of internal subunits with open backdoor gates was 0.6 in WT and 0.9 in N111S filaments, due entirely to the shorter sidechain of serine. Remarkably, the slopes of plots of gate separation vs. dihedral angle were similar for WT and N111S ADP-P_i_-actin filaments (Fig. 4B). The two lines are offset by about 1.5 Å at all dihedral angles owing to the shorter serine sidechain. This analysis did not detect any effect of the N111S mutation on the dihedral angle fluctuations that control the gate.

The N111S mutation decreased the probability of the occluded state. Figs. 4D-F document three differences in the interactions of R177 with its partners in the phosphate release channel in the N111S mutant actin: the probability of a hydrogen bond with the carbonyl oxygen of H73 is 20% lower (Fig. 4D); the probability of a hydrogen bond with the carbonyl oxygen of P109 unchanged (Fig 4E); and the probability of an electrostatic interaction with D179, which opens the channel, is two-fold higher (Fig. 4F), which likely accounts for the higher probability of R177 turning away from both H73 and P109 (Fig. 4D,E).

Reversible separation from R177 allowed the sidechain of N111 to rotate, which opened space for the sidechain of H161 to rotate from its gauche plus conformation into the gauche minus conformation. The gauche plus conformation is observed in cryo-EM structures of filaments with ATP analogues (Oosterheert, Klink et al. 2022, Reynolds, Hachicho et al. 2022) and a crystal structure of AMPPNP-actin bound to fragmin domain-1 in a filamentous conformation (Kanematsu, Narita et al. 2022, Iwasa, Takeda et al. 2023). During the simulations of ADP-P_i_-actin filaments, the average conformation of H161 in the internal subunits transitioned from gauche plus to gauche minus conformation five times faster in the N111S mutant actin (Fig. S8). The repositioning of the H161 side chain could affect the disruption of P_i_-Mg^2+^ interaction as noted in (Wang, Wu et al. 2024).

### Effects of subunit dynamics at 310 K on cofilin and phalloidin binding

Phalloidin, a fungal metabolite that stabilizes actin filaments and is conjugated to fluorescent dyes for use in many light microscopy studies, binds to actin filaments extremely slowly with an association rate constant of 0.03 μM^-1^s^-1^ for rhodamine-labelled phalloidin (De La Cruz and Pollard 1994). Phalloidin fills a cavity of ∼930 Å^2^ formed by three neighboring subunits where it is stabilized by hydrophobic interactions and hydrogen bonds (Fig. 5A) (Pospich, Merino and Raunser 2020).

**Figure 5:**
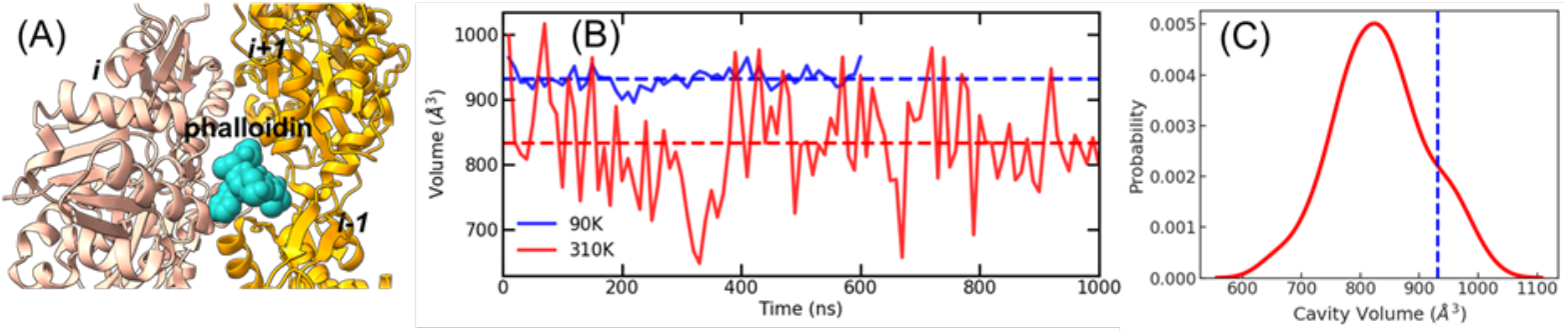
Phalloidin binding to ADP-actin filaments. (A) Ribbon diagram of the cryo-EM reconstruction of phalloidin bound among three subunits (pdb id: 6T20). (B) Time evolution of the volume of the phalloidin cavity formed by 3 adjacent internal subunits in 90 K and 310 K simulations of filaments with 27 ADP-actin subunits. Dashed line represents the average cavity volume. (C) Probability distribution of the phalloidin cavity volume at 310 K. The dotted vertical line represents the mean cavity volume at 90 K.

In 1 μs of MD simulations of the filament with 27 ADP-actin subunits at 310 K the volume of the phalloidin-binding cavities measured using POVME algorithm (Durrant, Votapka et al. 2014) varied from 620 Å^2^ to 1050 Å^2^ with a mean of 830 Å^2^, less than the volume occupied by phalloidin in the cryo-EM reconstruction pdb id: 6T20. Breathing motions of the filament during the simulation generated cavities larger than the volume that fits phalloidin only 14% of the time.

Cofilin binds between two subunits in ADP-actin filaments with an association rate constant of 0.03 μM^-1^s^-1^ (Blanchoin and Pollard 1999). Opening this binding site requires helical rotations of two consecutive subunits by between -161.4° to -162.4° along the short-pitch helix (pdb id: 5YU8(Tanaka, Takeda et al. 2018)). Single subunits sample these angles 9% of the time (Fig. 1D). Assuming that the adjacent subunits fluctuate independently, the binding site opens only 0.8% of the time. Surprisingly, the fraction of open sites is similar for ADP-P_i_ filaments, which is unexpected since cofilin has very low affinity for ADP-P_i_-actin (Blanchoin and Pollard 1999).

### Conformations of pointed end subunits of actin filaments during MD simulations at 310 K

Our new simulations of filaments with 27 or 7 ADP-actin subunits at 310 K confirmed the main findings our previous MD simulations of filaments with 13 subunits (Zsolnay, Katkar et al. 2020). Both subunits P and P-1 sampled wide ranges of conformations with mean dihedral angles more twisted than the starting cryo-EM structure (Fig. 1C, Figs. S1 and S9-12). Subunit P was more twisted in longer filaments (Fig. 6B Inset), perhaps due to finite size effects during the simulations. The salt bridge between R62 of subunit P-1 and E260 of subunit P formed as subunit P transitioned to a less twisted conformation in previous simulations of 13 subunit filaments (Zsolnay, Katkar et al. 2020). Although the transition to less twisted conformations occurs in filaments with 7 and 13 subunits in the new simulations, the less twisted conformations transition to more twisted conformations at longer times.

**Figure 6:**
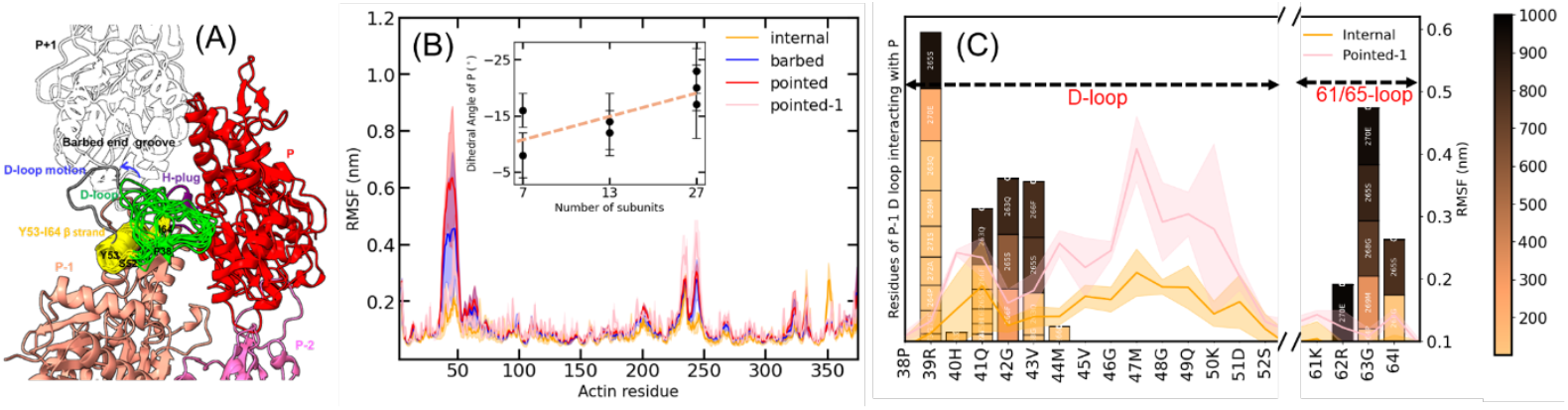
Conformations of subunits P and P-1 during simulations of ADP-actin filaments at 310 K in 1 μs with 27 subunits. (A) Ribbon diagram of subunits P (red), P-1 (coral pink) and P-2 (pink) at the end of the 1 μs MD simulation with P38 at the start and S52 at the end of D-loop labelled. The D-loop of P-1 (green) is rendered every 100 ns to show the persistent, variable interactions with the H-plug (purple) of subunit. P that prevent the D-loop from interacting with an incoming monomer P+1 (black outline), which requires an extended D-loop (grey backbone, blue arrow) to bind the barbed end groove of subunit P+1 as in internal subunits. (B) Comparison of the root mean square fluctuations (RMSF) of each residue of internal (yellow), B (blue), P (red) and P-1 (pink) subunits. RMSF of D-loop residues are P > B > P-1 and I. (Inset) Average dihedral angles (±SD) of subunit P in replica simulations of filaments with 7, 13 and 27 subunits. (C) Interactions of residues in SD2 of subunit P-1 with the H-plug of subunit P that are within 4.5 Å for more than 400 ns. Color bars and heights of the boxes represent total times that P residues contacted H-plug residues of P-1 identified by white numbers in the boxes. Heights of the bars represent promiscuity of the interactions. D-loop and flanking residues R39 and Q41-V43 and loop I64-G63 residues contacts with the H-plug persistently. Solid line traces are the RMSFs of residues on SD2 of internal (yellow) and P-1 (pink) subunits.

The D-loop (residues R39 to S52) of subunit P did not interact with neighboring subunits and sampled many conformations as indicated by the high RMSF (Fig. 6B and Movie S1). Bending of SD2 relative to SD1 contributed more to the twisted conformation of subunit P than rotation between outer domain and inner domain in monomers (Fig. S9 D-F; Table S1).

Our new simulations confirmed that the P-1 D-loop bent towards subunit P and interacted its H-plug (residues Q263 to A272) (Fig. 6A), as observed our previous simulations of a 13-mer actin filament (Zsolnay, Katkar et al. 2020). Figure 6C compares the RMSD fluctuations of D-loop residues with the frequency of interactions with the H-plug of subunit P. The residues R39 to M44 maintained persistent contacts with the H-plug of subunit P (Fig. 6C), while. exploring many conformations (Fig. 6A (green) and Movie S1). These contacts between P-1 D-loop residues (R39, Q41, G42, V43) and subunit P H-plug residues R62, G63, I64 depend on SD2 of P-1 bending toward subunit P, which contributes to the twisted conformation of subunit P-1. The interactions between D-loop of P-1 and P would limit interactions of the D-loop with an incoming monomer P+1, which requires an extended D-loop (grey in Fig. 6A). Subunit P-1 D-loop residues V45 to S52 had high per-residue root mean square fluctuations and no contacts with the P subunit (Fig. 6C). Residues R62 to I64 in the loop before the final ß-strand in SD2 form a secondary contact with subunit P (Fig. 6C).

### Conformations of barbed end subunits of actin filaments during MD simulations at 310 K

In cryo-EM reconstructions of barbed ends (pdb id: 8F8R) (Carman, Barrie et al. 2023) subunits B and B-1 are flattened like internal subunits, but they were dynamic and more twisted in our new MD simulations of filaments with 27 or 7 subunits as in our simulations of filaments with 13 subunits (Zsolnay, Katkar et al. 2020), especially when subunit B dissociated laterally from subunit B-1 (Figs. 7 and S13, Movies S2 and S3). Subunit B was stably tethered to subunit B-2 but lost and reformed lateral contacts with subunit B-1 (Figs. 7 and S13 and Movies S2 and S3). Like internal subunits (Fig. 4D yellow lines) the longitudinal anchor between the D-loop of subunit B and barbed end groove of subunit B-2 persisted throughout all the simulations covering 6.4 μs (Movies S1 and S3, Figs. 4D and S10A three blue lines). Other longitudinal contacts in the starting structure between SD4 residues I287-K291 and A321-K326 of subunit B and SD3 residues P243-Q246 of subunit B-2 (referred to here as SD3/SD4 contacts) were lost over time (Figs. 7E and S13A blue lines), but were stable between internal subunits (Fig. 7E and S13A yellow lines). The lateral contacts between SD4 of subunit B with SD1 of subunit B-1 were unstable, dissociating and reforming randomly in nine simulations including filaments with 7, 13 or 27 subunits (Figs. 7A-C, F and S13A). In the example in Fig. 4F and Movie S1 the lateral contacts weakened around 350 ns and reformed by the end of the simulation, while in the third replica simulation the lateral contact dissociated and reformed more often (Fig. S13A and Movie S2 and S3).

**Figure 7:**
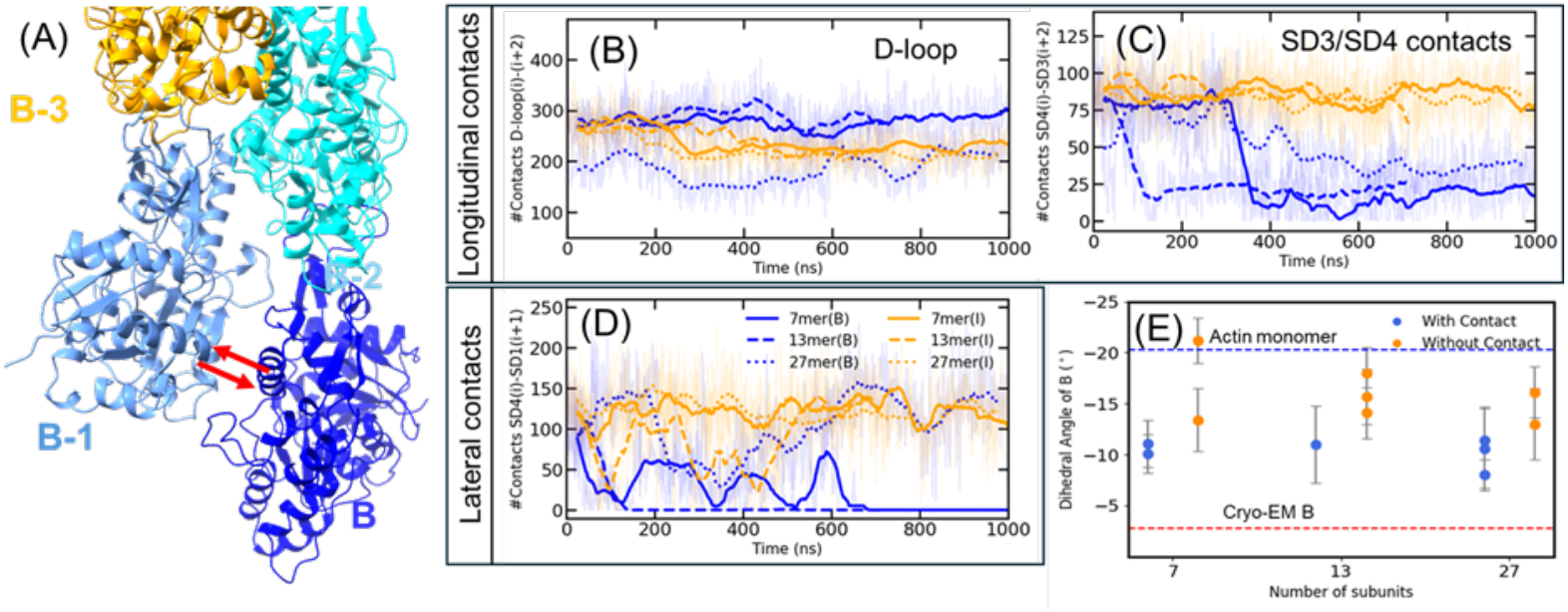
Conformations and contacts of internal subunits and barbed end subunit B during MD simulations. (A) Ribbon diagrams of the barbed end at time point ∼350 ns in the simulation of the 27-mer when subunit B was dissociated from B-1 while tethered to B-2. Red arrows indicate reversible association of subunit B. (B-E) Time evolution of the number of contacts within a cut-off of 3.5 Å for internal subunits (I) and terminal subunit B in filaments with 7, 13 or 27 subunits. Longitudinal contacts with the next subunit along the long-pitch helix were calculated in two separate groups: (B) D-loop contacts made by SD2 of subunit B were stable like the interactions between internal subunits; and (C) other contacts made by SD4 with SD3 were unstable for subunit B but stable for internal subunits. (D) Lateral contacts between SD4 of a subunit and SD1 of the subunit across the filament up the short-pitch helix. Subunit B dissociated reversibly from subunit B-1 during the simulations of all three filaments, while the lateral contacts between internal subunits are stable. (E) Dihedral angles (mean ± SD) of barbed subunit B in actin filaments with 7, 13 and 27 subunits during segments of the simulation (Table S3) with and without lateral contacts. Dashed red line is the dihedral angle of subunit B in the cryo-EM reconstruction, and dashed blue line is the dihedral angle of an actin monomer.

The B subunits were more twisted when they lost their lateral and SD3/SD4 contacts than in filaments with these contacts and had more negative dihedral angles than the starting structure at all filament lengths (Fig. 7G using the data in Table S3). Another consequence of the loss of lateral and longitudinal contacts and twisting of subunit B was larger fluctuations in the center-of-mass of subunit B especially in shorter filaments (Fig S13B).

## Discussion

Samples with monodisperse structurally homogeneous particles are ideal for obtaining invaluable high-resolution reconstructions with a good signal to noise ratio from cryo-EM images (Cheng 2015, Cheng, Grigorieff et al. 2015). Algorithmic developments bolstered by advances in machine learning methods can recover conformational heterogeneity in cryo-EM images (Nakane, Kimanius et al. 2018, Luo, Ni et al. 2023, Punjani and Fleet 2023, Tang, Zhong et al. 2023).

However, proteins and protein assemblies can have a different Boltzmann distribution of conformations at physiological temperatures that may be lost during freezing. Preparation for single-particle cryo-EM imaging involves depositing a thin film of hydrated particles on an EM grid and plunge freezing the sample in liquid ethane at *T* ≃ 90*K* or ethane/propane mixture at *T* ≃ 77*K*. Plunge cooling rates of the order 10^5^ − 10^8^*K*/*s*, corresponding to upper timescales of 2-3 milliseconds, are used to avoid crystallization of water (Dubochet, Adrian et al. 1988, Engstrom, Clinger et al. 2021). It is assumed that these “fast” cooling rates of macromolecules trap the conformations at physiological and laboratory temperatures prior to freezing. However, computational and theoretical studies have shown that protein sidechains and conformations can change during the time required for freezing (Nagai, Tama and Miyashita 2019, Bock and Grubmüller 2022, Bock, Igaev and Grubmüller 2024). During cooling, barriers between 2.0 -2.5 kcal/mol are estimated to possibly be crossed while the molecule equilibrates to lower free energy conformations (Bock and Grubmüller 2022).

Molecular Dynamics with Flexible Fitting (Trabuco, Villa et al. 2008) (MD-FF) has been used to refine cryo-EM reconstructions of actin filaments at temperatures below 120 K (Reynolds, Hachicho et al. 2022), which did not change the subunit conformation. Cryo-EM models of other proteins can change during MD simulations at higher temperatures. For example, during MD simulations at 200 K, starting from of a cryo-EM reconstruction of the Ca^2+^ ATPase pump the conformations of some sidechains changed to agree with physiology experiments (Miyashita, Kobayashi et al. 2017). These sidechains in both the original model and the refined model fit equally well into the density map. MD simulations of hERG ion channels at 310 K revealed functionally relevant interactions that were not present in the starting cryo-EM reconstruction (Khan, Guo et al. 2021).

Here, we investigated how being frozen affects the structure of a multi-protein complex using actin filaments as our scientifically important example. We find that actin filament subunits adopt a lower entropy state in frozen samples that, when projected to physiological temperature, differs from the ensemble of conformations at higher temperature (and one which satisfies the Principle of Maximum Entropy). We then report the many consequences of this result as follows.

### Lower entropy ordered conformations of cryo-EM samples

Thermal fluctuations at room temperature confer flexibility to actin filaments that allow the subunits to adopt a collection of higher entropy conformations, including more twisted subunits with a range of dihedral and helical rotation angles. On the other hand, the frozen filaments used for cryo-EM imaging are homogeneous enough to make high resolution reconstructions with fixed helical rotations and subunit dihedral angles of lower entropy, even when the covariance matrix of a “flat” conformation is scaled to 310 K. Our entropy calculations of these actin subunits at 310 K showed that the twisted subunits have four times higher entropy than flat subunits, which explains the conformation after freezing. This analysis shows that the all-atom MD data are critical for understanding differences between filament structures at 310 K and cryo-EM structures at 90 K.

### Internal subunits are more twisted at 310 K

During atomistic MD simulations of actin filaments at 310 K, internal subunits sampled a wide range of dihedral angles between -17° to -3° with a mean value 9° more twisted than the starting cryo-EM reconstruction (Fig. 1D). The absence of free energy barriers separating these conformations (Fig. 2F) explains how this ensemble of conformations exists at 310 K. Transient losses of lateral contacts between subunits due to thermal fluctuations facilitates the wide variation in the dihedral angles of the internal subunits (Movie S1). On the other hand, flatter subunit conformations are enthalpically preferred at lower temperatures, so they predominate in cryo-EM reconstructions of internal actin subunits.

### Insights about ATP hydrolysis

The most detailed information about the active site in ATP-actin filaments comes from high resolution structures assumed to be good analogues of the Mg^2+^-ATP-actin filament: a 2.2 Å resolution cryo-EM reconstruction of the internal subunits in the Mg^2+^-ADP-BeF_3_-actin filament (pdb id: 8A2R; (8)) and 1.15 Å resolution frozen crystal structure of an AMPPNP-actin monomer held in a filamentous conformation by fragmin domain-1 (pdb id: 7W4Z) (Kanematsu, Narita et al. 2022, Iwasa, Takeda et al. 2023). In both structures, OE1 of Q137 coordinates a water molecule (W1) for in-line attack on the γ-phosphate of ATP for hydrolysis (Fig. 3A) (Oosterheert, Klink et al. 2022). Residues H161 and D157 position through hydrogen bonds a second water (W2) to activate the catalytic water. The geometries are close to ideal for catalysing ATP hydrolysis, but the rate of ATP hydrolysis by Mg-ATP-actin filaments, 0.3 s^-1^ (Blanchoin and Pollard 2002), is orders of magnitude lower than expected for an enzyme with a prebound substrate and the catalytic water positioned by the sidechain of Q137 to attack the γ-phosphate of ATP.

Prior studies (McCullagh, Saunders and Voth 2014, Sun, Sode et al. 2017) account for the dynamic conformations at 310 K in determining the free energy barrier of ATP hydrolysis using QM/MM MD simulations with metadynamics to enhance the sampling. The QM/MM minimum energy reaction pathway studies in Kanematsu et al. (Kanematsu, Narita et al. 2022) provide a zero temperature potential energy barrier to ATP hydrolysis. The conformations of the active site residues and positions of water molecules sampled at 310 K in this work and that of (McCullagh, Saunders and Voth 2014, Sun, Sode et al. 2017) incorporate the important thermal fluctuations, which reduce the probability of conformations that favor ATP hydrolysis in actin monomers and actin filament subunits.

Our MD simulations at 310 K captured molecular motions in the active site that include conformations favorable for ATP hydrolysis, but these conformations occur only a small fraction of the time. At 310 K, the conformation of the sidechain of Q137, the angle of attack, distance and orientation of the catalytic water 1 and water 2 positioned to activate water 1 are favorable less than 0.00012 of the time in interior subunits of Mg-ATP actin filaments (Fig. 3 and associated text). Thus, the overall rate of hydrolysis, 0.3 s^-1^, is more than 8000 times lower than the rate of 2500 s^-1^ in rare but suitably configured active sites of subunits in the filament. This is consistent with work on thermal fluctuations of side chains and water in the active sites of enzymes at 310 K, which allow them to sample numerous states not observed in cryo-EM reconstructions (Wankowicz and Fraser 2025).

This requirement for thermal motions to optimize the active site was suggested by the high resolution structures of the active site (8, 36, 37). Oosterheert et al. (8) noted the catalytic water “probably exchanges between hydrolysis-competent and hydrolysis-less-competent configurations” (in their cryo-EM reconstruction). Katematsu et al. (Kanematsu, Narita et al. 2022, Iwasa, Takeda et al. 2023) observed “The W2 fluctuations (detected by the shape of the electron density) may partly explain the relatively slow rate of actin-mediated ATP hydrolysis.”

The active site of an actin monomer visits ideal catalytic conformations so rarely that they were not detected during 4 μs of simulations. This is consistent with ATP-actin monomers hydrolyzing ATP 42,000 times slower than subunits of Mg-ATP actin filaments (Blanchoin and Pollard 1999, Rould, Wan et al. 2006, McCullagh, Saunders and Voth 2014, Sun, Sode et al. 2017).

### Insights about dissociation of the γ-phosphate of ATP

In contrast to most cryo-EM reconstructions with closed backdoor gates between N111 and R177 blocking the escape of the γ-phosphate of ATP (Merino, Pospich et al. 2018, Oosterheert, Klink et al. 2022), the gate is open about 60% of the time for internal subunits in our MD simulations of ADP-P_i_ actin filaments at 310 K (Fig. 4). The distance between N111 and R177 is coupled to the dihedral angle of the subunit during the thermal motions of the subunits, because these two gate residues are in different halves of the molecule (Fig. 4A). These open gates cannot, however, explain the very slow rate of phosphate release. The large energy barrier associated with dissociation of phosphate from Mg^2+^ makes it the slow step (Wriggers and Schulten 1999, Wang, Wu et al. 2024).

We simulated the ADP-P_i_ actin filament with the mutation N111S at 310 K to distinguish between two hypotheses to explain how the mutation increases the rate of phosphate dissociation 15-fold (Oosterheert, Blanc et al. 2023). The authors of the original paper (Oosterheert, Blanc et al. 2023) argued that the mutation increases the probability that the backdoor gate is open. Further, they concluded (Oosterheert, Boiero Sanders et al. 2025) that their mutagenesis experiment is inconsistent with enhanced sampling MD simulations showing that dissociation phosphate from Mg^2+^ is rate limiting step (Wang, Wu et al. 2024). Our new simulations show that the N111S mutation increases the fraction of open gates modestly but also decreases the probability of occluded states. (In addition, the N111S mutation alters the dynamics of H161 flipping from gauche plus to gauche minus state, which may favor to some degree the escape of phosphate from Mg^2+^ (Fig. S8).) Furthermore, it is important to recognize that a 15-fold increase in the rate of P_i_ release by N111S actin amounts to only a 2 kcal/mol reduction in the overall kinetic energy barrier of ∼ 20 kcal/mol that determines the long timescale of P_i_ release.

### Breathing motions of the ADP-actin filament explain slow binding of cofilin and phalloidin

Both the protein cofilin (Blanchoin and Pollard 1999) and the small fungal metabolite phalloidin (De La Cruz and Pollard 1994) bind very slowly to ADP-actin filaments (De La Cruz and Pollard 1994), compared with expectations of the Debye-Smoluchowski equation for diffusion-limited collisions (Berg and von Hippel 1985). Our MD simulations show that breathing motions of the filament arising from thermal fluctuations in the dihedral angles and helical rotation angles open up both binding sites that are not accessible in cryo-EM reconstructions of filaments (Carman, Barrie et al. 2023, Chou and Pollard 2023, Oosterheert, Blanc et al. 2023, Boiero Sanders, Oosterheert et al. 2024, Oosterheert, Boiero Sanders et al. 2024).

Rhodamine-phalloidin binds between three subunits in ADP-actin filaments with a very small association rate constant of 0.0003 μM^-1^s^-1^ (De La Cruz and Pollard 1994). The rhodamine conjugated to phalloidin probably protrudes from the filament surface. Our MD simulations show that a volume sufficient to bind phalloidin opens up 14% of the time (Fig. 7D). Correcting for the fraction of sites large enough to bind phalloidin gives an association rate constants of 0.002 μM^-1^s^-1^, far less than 100 μM^-1^s^-1^, the rate constant for collisions expected from the Debye-Smoluchowski equation. Therefore, rhodamine conjugated to phalloidin may compromise binding or the orientation factor must be very small, as one might expect from the narrow opening of the flat binding site.

Cofilin binds between two subunits in ADP-actin filaments with an association rate constant of 0.03 μM^-1^s^-1^ (Blanchoin and Pollard 1999). Assuming that the adjacent subunits fluctuate independently, our MD simulations show that the binding site opens only 0.8% of the time. Taking into account the fraction of available sites, the association rate constant for cofilin is ∼ 4 μM^-1^s^-1^, close to the Debye-Smoluchowski collision rate corrected by a relatively large orientation factor around 0.04, so binding is very favorable. Concerted rotations of consecutive subunits might make this breathing more or less favorable for cofilin binding.

However, much remains to be learned about cofilin binding to actin filaments, since our simulations did not reveal why cofilin binds extremely weakly to ADP-P_i_-actin filaments (Fig. 1D). In addition to the fluctuations in the short-pitch helix which open up the binding site (Blanchoin and Pollard 1999); and fluctuations of SD1 may open up a partial binding site for cofilin binding followed further changes through induced fit to complete binding (Tanaka, Takeda et al. 2018). Phosphate in the active site may influence this induced fit process.

### Behaviour of barbed end subunits at 310 K

During our MD simulations of filaments with 7, 13 and 27 subunits at 310 K, subunit B was persistently tethered to subunit B-2 by longitudinal interactions mediated by its D-loop but stochastically lost lateral contacts with subunit B-1 and longitudinal SD3/SD4 contacts with subunit B-2 (Fig. 3). These dissociation events were more common in simulations of shorter filaments. When subunit B dissociated from B-1, both subunits were more twisted that the cryo-EM reconstruction (PDB ID: 8F8R).

These observations on barbed ends support the hypothesis based on MD simulations of Zsolnay et al. (Zsolnay, Katkar et al. 2020) and cryo-EM structures of Palmer et al.(Palmer, Barrie and Dominguez 2024) that addition of an actin monomer to a barbed end begins with its D-loop binding to an open hydrophobic pocket above the W-loop of subunit B-1. This longitudinal interaction of incoming subunit B+1 is more favorable than binding laterally to the loosely tethered and often twisted subunit B. Subsequent formation of lateral contacts between tethered subunit B+1 with subunit B stabilize the new subunit and favor flatter conformations of subunit B.

### Behavior of pointed end subunits

Our new MD simulations of filaments with 7 and 27 subunits confirmed and extended MD simulations on filaments with 13 subunits (Zsolnay, Katkar et al. 2020). They show that subunits P and P-1 are more dynamic than internal subunits (Figs. 1CD, 2AB) with large variances in the dihedral angles (Fig. 1C) and broader free energy basins than internal subunits and subunit B (Fig. 4D-E). Like the twisted pointed end subunits in the cryo-EM reconstruction (pdb id: 8F8S) (Carman, Barrie et al. 2023), subunits P and P-1 are more twisted than the internal subunits during our MD simulations owing to bending between SD1 and SD2, rather twisting between SD2 and SD3.

The viscosity dependence of the subunit association rate constant at the pointed end demonstrated that the reaction is not diffusion limited (Drenckhahn and Pollard 1986). This is explained by the time required to sample conformational rearrangements that overcome two features that slow subunit addition, limited availability of the D-loop of P-1 to bind incoming subunit P+1 and steric hinderance between the D-loop of twisted subunit P and incoming subunit P+1. Interactions of P-1 with P may also contribute to the very slow rate of dissociation of ADP-actin from the pointed end, 0.27 s^-1^ (Pollard 1986).

### Concluding Remarks

Our study shows that a low entropy sub-population of conformations of actin filaments become favoured as the sample cools during freezing and dominates in cryo-EM samples. However, at physiological temperature this sub-population is significantly less probable than the collection of populations corresponding to higher entropy. Our observations on actin filaments demonstrate the value of applying physics-based MD simulations at 310 K to cryo-EM-based structural models, to reveal the operational structures and dynamics at physiological temperature.

## Methods

### Simulation set-up

The 27 subunit actin filament was constructed by stitching together the barbed (pdb id: 8F8R), internal (pdb id: 8F8P) and pointed (pdb id: 8F8S) end cryo-EM structures (Carman, Barrie et al. 2023). The terminal pointed end subunit in the cryo-EM structure lacks the coordinates for ten residues (41-51) in the D-loop. These 10 residues were modelled using Charmm-GUI (Jo, Kim et al. 2008). The filament was placed in a cubic periodic box with a minimum distance of 1.2 nm between the protein and the box boundary. The protein was solvated using TIP3P water molecules (Jorgensen, Chandrasekhar et al. 1983). KCl concentration was set to 100 mM at which the system was in charge-neutral state. The total system size was 1.3 million atoms for the 27 subunits filament. The simulations are done using the CHARMM36 force field (Huang, Rauscher et al. 2017). Similar protocol was followed to set up simulations of filaments of length 7 subunits (342,000 atoms) and 13 subunits (692,000 atoms).

### MD simulations

Electrostatics were treated using particle mesh Ewald sum method with a cut-off of 12 Å. A cutoff distance of 12 Å was used for the nonbonded interactions. Bond lengths involving hydrogens were constrained using the LINCS algorithm (Hess, Bekker et al. 1997). The solvated filament was first energy minimized with harmonic restrains of 400 kJ/mol on the backbone atoms and 40 kJ/mol on the side chain atoms and LINCS constraints on the H-bonds. Following the energy minimization, the restrains on the backbone and side chain atoms were sequentially removed while the solvated protein equilibrated for 25 ns to the set temperature of 310 K and pressure of 1 bar. Following the restrained equilibration, an unconstrained equilibration for 12 ns was performed in the constant NPT ensemble using a Nose-Hoover thermostat with a 1.0 ps coupling time constant and Berendsen barostat with a 5.0 ps coupling time constant (Berendsen, Postma et al. 1984). The simulation was then continued at constant NPT for 1 μs of production run with a 2 fs timestep. Pressure in the production was regulated using the Parrinello-Rahman barostat with a coupling constant of 5.0 ps (Parrinello and Rahman 1981). Three replica simulations were run for the 27-mer ADP filament for 1 μs,450 ns and 450 ns respectively. The wild type and (2 replica) mutant ADP-Pi 27 subunit filament simulations (constructed from pdb id: 8A2R) were run for 400 ns and 300 ns respectively. Three replica simulations of lengths 1.17 μs, 1.04μs and 380 ns respectively were run for seven subunit filaments. Three replica simulations of lengths 740 ns, 730 ns and 430 ns respectively were run for thirteen subunit filaments. The results for the replica runs are presented in the supporting information. Additionally, a 13-mer ADP actin filament was simulated at five other temperatures: 175 K, 230 K, 250 K, 275 K and 298 K for 330 ns, 280 ns, 580 ns and 250 ns respectively.

### Enhanced sampling using well-tempered metadynamics

The two-dimensional free energy of subunit twisting was calculated using well-tempered metadynamics with dihedral angle and cleft width as two collective variables. The free energy of twisting was calculated for the terminal and internal actin subunits in a filament of 7 actin subunits. The height of Gaussian bias was set to 0.06 kcal/mol and a biasing potential was added every 1000MD steps. The biasing factor was set to 6 and gaussian widths of 2° and 1 Å for dihedral angle and distance respectively, was used for all the simulations. The convergence of the simulation was accessed based on the difference in free energy projected on the dihedral angle collective variable (Fig. S12). All the MD simulations were preformed using GROMACS 2021.5 (Abraham, Murtola et al. 2015) with PLUMED 2.7 (Bonomi, Branduardi et al. 2009).

### Entropy calculation

Schlitter’s method of entropy calculation using the covariance matrix of atom-positional fluctuations is used to estimate backbone conformational entropy of the actin subunits (Schlitter 1993). Using the quasi-harmonic-approximation, entropy is given as:

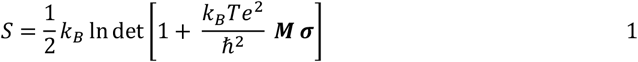

*k*_*B*_ is Boltzmann’s constant, *T* is the absolute temperature, *e* Euler’s number, *ℏ* is Planck’s constant divided by 2π, ***M*** is 3N dimensional the mass matrix and ***σ*** covariance matrix of atom-positional fluctuations. Its elements are given by:

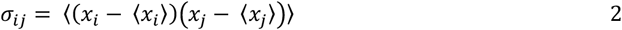

The averaging, indicated by ⟨·⟩, is performed over all trajectory frames up to a certain time to yield the entropy buildup overtime. The initial configuration after equilibration was used as a reference structure for a least-squares fit of trajectory configurations to remove overall translational and rotational motion. The entropy of different average dihedral twist conformations at 310 K is calculated by scaling the fluctuations measured at the respective temperatures to 310 K. That is, the correlation matrix ***M*** was measured for average dihedral angles sampled 90 K, 175 K, 230 K, 250 K, 270 K, 298 K and 310 K, and to calculate the entropy of the corresponding average dihedral angle conformations, the temperature in Eqn (1) was set to 310 K.

### Analysis

The dihedral angle of actin subunits was calculated based on the center of mass of the Cα atoms of each subdomain. The subdomains are defined as subdomain 1 (SD1): residues 1 to 32, 70 to 144, and 338 to 375; (SD2): residue 33 to 69; (SD3): residue 145 to 180, 270 to 337; (SD4): residue 181 to 269.

### Data, Materials, and Software Availability

The study data are available upon request due to large file sizes and number of files.

## ACKNOWLEDGMENTS

Research reported in this publication was supported by the National Institute of General Medical Sciences of the NIH under award number R01GM063796 to G.A.V. The content is solely the responsibility of the authors and does not necessarily represent the official views of the NIH. Computational resources were provided by the University of Chicago Research Computing Center and the NIH-funded Beagle-3 computer (NIH award 1S10OD028655-01).

