## Supporting Information for "Reaching the full potential of cryo-EM reconstructions with molecular dynamics simulations at 310 K: Actin filaments as an example"

#### Section 1: Replica MD simulations of an ADP-actin filament with 27 subunits

Analysis of the 450 ns simulations in second and third replica trajectories of the of 27-mer actin filament at 310 K, which agree with the first replica results reported in Fig. 1.

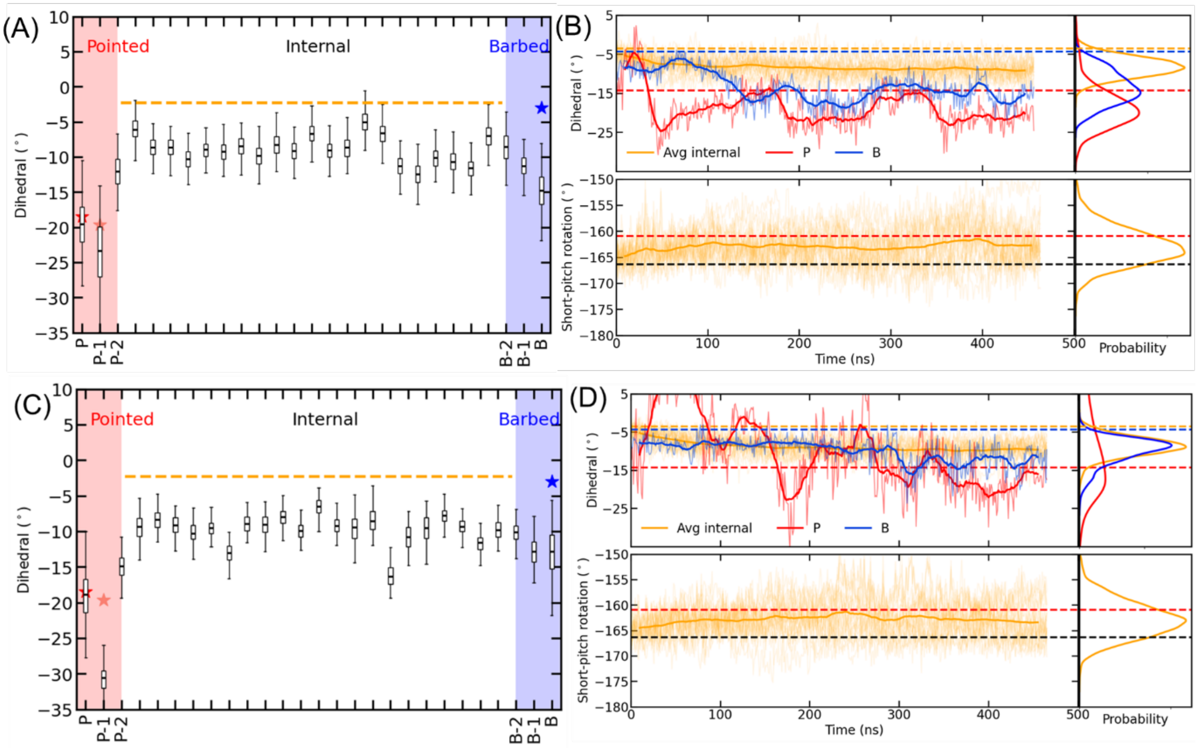


**Figure S1**: Simulations of a 27-subunit ADP-actin filament for 450 ns in (A, B) the second replica and (C,D) the third replica. (A, C) Distributions of dihedral angle of all subunits during the last 200 ns of replicas 2 and 3. Edges of the box plots represent first and third quartiles and the horizontal lines within each box are mean dihedral values. Whiskers extend to minimum and maximum dihedral angles sampled. (B, D) (Top) Time course of dihedral angles of the terminal barbed end subunits B, pointed end subunits P, and average of the 23 internal subunits during the MD simulations. (Bottom) Time courses of helical rotation angles averaged across all internal subunits. Dashed lines indicate the dihedral and helical rotation angles of the subunits in the cryo-EM reconstructions. Darker lines are running averages and the lighter lines are the raw data from the simulations.

**Summary of subunit dihedral angles sampled in MD simulations and cryo-EM reconstructed structures**

**Table S1:** Summary of subunit dihedral angles as defined in Fig. 1B in cryo-EM reconstructions and during the last 200 ns of the three MD simulations of the ADP-actin filament with 27 subunits (average ±SD). Dihedral angles were calculated for subunit P with and without the D-loop to show that the bending of D-loop of P subunit, rather than the rotation of inner subdomains generates subunit twisting.

| Subunit | Carman cryo-EM reconstructions | MD at 310K (last 200 ns) Rep1, (Rep2,Rep3) | | MD at 90 K (last 200 ns) |
|---|---|---|---|---|
| | | With D loop | w/o D loop | |
| P | -18.5 (no D-loop) PBD 8F8S | -23.2 ±4.0 (-19.7 ±3.7,-15.5±6.5) | -19.6 ±3.9 (-15.8 ±3.6,- 13.0±4.5) | -13.7 ±0.3 (w/o D loop) |
| P-1 | -19.6 (no D loop) PDB 8F8S | -22.3 ±1.6 (-23.3 ±4.4,-30 ±1.5) | -22.4 ±1.6 (-19.8 ±4.0,-22.1±1.7) | -18.2 ±0.3 (w/o D loop) |
| P-2 | -2.2 PDB 8F8S | -9.0 ±1.6 (-12.0±2.3, -17.2 ±2.0) | -9.3 ±1.5 (-12.2 ±1.9,-16.2±2.1) | -2.2 ±0.2 (with D loop) |
| Internal | PDB 8F8P -2.3 | -9.1±2.4 (-9.3 ±2.2, -9.6±2.5) | -9.1±2.4 (-8.9 ±2.5, -9.9±2.3) | -2.3±0.6 (with D loop) |
| B | -3.0 PDB 8F8R | -12.8 ±2.6 (-14.8±2.8, -13.7 ±3.7) | -14.6 ±2.4 (-15.2 ±2.6,-13.1±3.3) | -4.2 ±0.2 (with D loop) |
| B-1 | -2.6 PDB 8F8R | -9.6 ±2.3 (-11.2 ±1.8,-13.8 ±2.4) | -10.6 ±2.2 (-11.6 ±1.6,-14.9±2.0) | -3.0 ± 0.2 (with D loop) |
| B-2 | -2.3 PDB 8F8R | -6.8 ±1.3 (-8.6±2.4, -11.5±1.6) | -8.0 ±1.3 (-9.6 ±2.0-11.0±1.5) | -2.2 ± 0.2 (with D loop) |

**Summary of short-pitch helix angles sampled in MD simulations and cryo-EM reconstructed structures**

**Table S2:** Summary of short-pitch rotation angles in cryo-EM reconstructions and during the last 200 ns of the three MD simulations of the ADP-actin filament with 27 subunits (average ±SD).

| Subunit | Carman cryo-EM reconstructions | MD at 310K (last 200 ns) Rep1, (Rep2,Rep3) | MD at 90K (last 200 ns) |
|---|---|---|---|
| P-1 | -166.3 PDB 8F8S | $-163.2 \pm 5.2$ ($-148.3\pm6.7$, $-158.4 \pm5.2$) | $-166.3\pm0.2$ |
| P-2 | -166.1 PDB 8F8S | $-158.5 \pm3.8$ ($-147.5\pm6.0$, $-154.6\pm5.2$) | $-166.6 \pm0.1$ |
| Internal | -166.8 to -166.5 PDB 8F8P | $-162.6 \pm 3.8$ ($-162.7 \pm3.5$, $-162.8\pm3.5$) | $-166.6\pm0.4$ |
| B | -166.1 PDB 8F8R | $-168.5 \pm4.4$ ($-155.2\pm8.1$, $-161.3 \pm4.6$) | $-166.0 \pm0.1$ |
| B-1 | -166.6 PDB 8F8R | $-164.3 \pm3.6$ ($-162.1 \pm4.6$, $-160.6\pm4.8$) | $-166.9 \pm 0.1$ |
| B-2 | -166.7 PDB 8F8R | $-162.7 \pm3.6$ ($-169.6\pm3.8$, $-159.5\pm3.7$) | $-166.5 \pm 0.2$ |

**Section 2: Persistence length calculation**

Persistence length ($L_p$) measure the stiffness of actin filaments. It is the length over which the direction of the filament remains correlated due to thermal fluctuations. The persistence length of ADP-actin filaments in solution at 293 K was measured as 9±0.5 μm (1).

We followed the procedure outlined in Chu and Voth (2) to compute the persistence length from the all-atom simulation of 27-mer and 13-mer ADP-actin filaments. The double helical filament was first mapped onto a one-dimensional linear polymer. The center of mass of each subunit ($i$) in strands was calculated. The centres of mass of adjacent subunits *i-1* and *i+1* was used to define $i'$. The positions were calculated every 1 ns in the last 500 ns of the 1 μs simulation. The midpoint of *i* and $i'$ represented the position of the $r_i$ monomer on the mapped linear polymer. The contour length was the distance between $r_i$ and $r_{i+1}$. Tangent vectors were computed from these positions and angle between consecutive tangent vectors was measured to give the bending angle. The persistence length was computed from the slope of the exponential decay of ensemble averaged cosine of the bending angle fluctuations as a function of the contour length:

$$\langle \cos\left(\theta(s)\right\rangle = exp\left(-\frac{s}{L_p}\right)$$

The persistence length calculated for the ADP-actin subunit filament with 27 subunits was 8 ±0.5 µm, close to the experimentally measured value of 9 ±0.5 µm. On the other hand, the cosine bending angle fluctuations for the longer contour length in 13 subunit ADP-actin filament deviated from the exponential decay. A longer filament of 27 subunits thus captures the experimentally observed persistence length, while shorter filament simulations show deviations due to finite size effects (3).

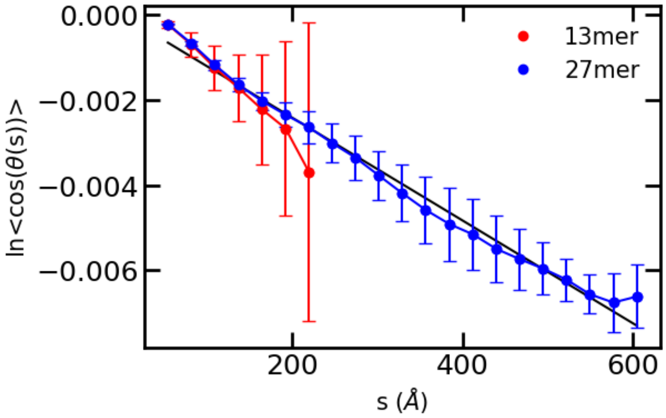


**Figure S2:** Persistence length calculation: Log of the cosine of bending angles of ADP-actin filaments with 27 and 13-subunits as a function of contour length. Error bars are SD measured by splitting the trajectory into blocks of 100 ns.

**Section 3: MD simulations of an ADP-actin filament with 13 subunits at 175 K, 230 K, 273 K, 298 K and 310 K.**

MD simulations of ADP-actin filaments was performed at 175K, 230K, 273K, 298K and 310K to study temperature dependence of the conformations of internal subunits.

Subunit dihedral angle: The average dihedral angle calculated across 8 internal subunits in the 13-mer (Fig. S9) indicates that average dihedral angle of the internal subunits decreases monotonically with temperature. i.e. it becomes more twisted at higher temperatures. The fluctuations in the dihedral angles (given by the SD) increases with temperature. Convergence of the simulations at different temperatures is monitored by plateauing of the average internal subunit dihedral angle (Fig. S9).

Helical rotation angle: The helical rotation angle measured between i and i+2 internal subunits increases monotonically with temperature (Fig. S10). The average and mean helical rotation angle (averaged across the 8 internal subunits) is similar to angle measured in the cryo-EM reconstructed F-actin structures below 230K. The fluctuations in the helical rotation angle (given by the SD in Fig. S10 labels and width of the probability distribution) increase with temperature.

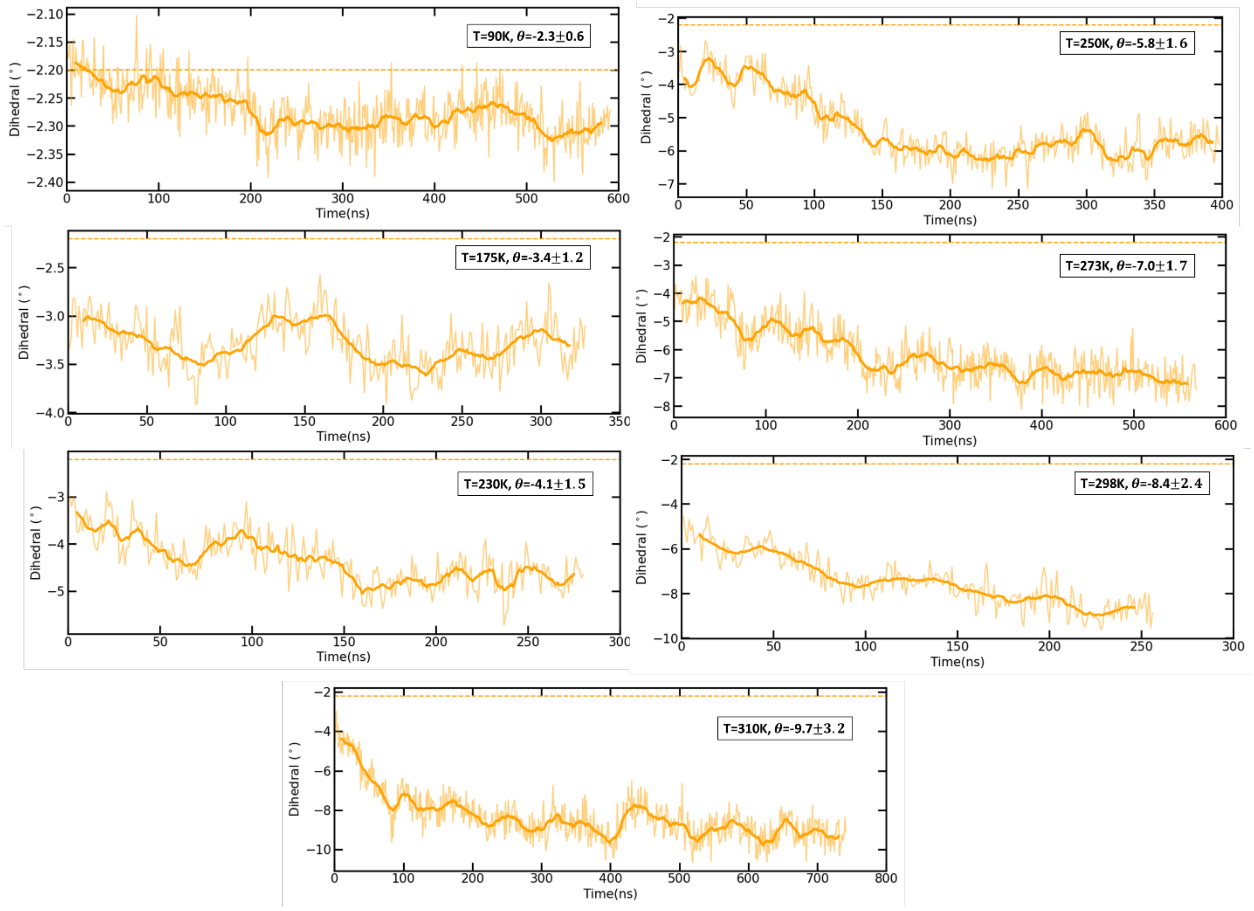


**Figure S3:** Time evolution of average dihedral angles of 7 internal subunits in 13-mer ADP-actin filaments at different temperatures. Text inserts are the averages and SDs during the last 100 ns of the equilibrated trajectory.

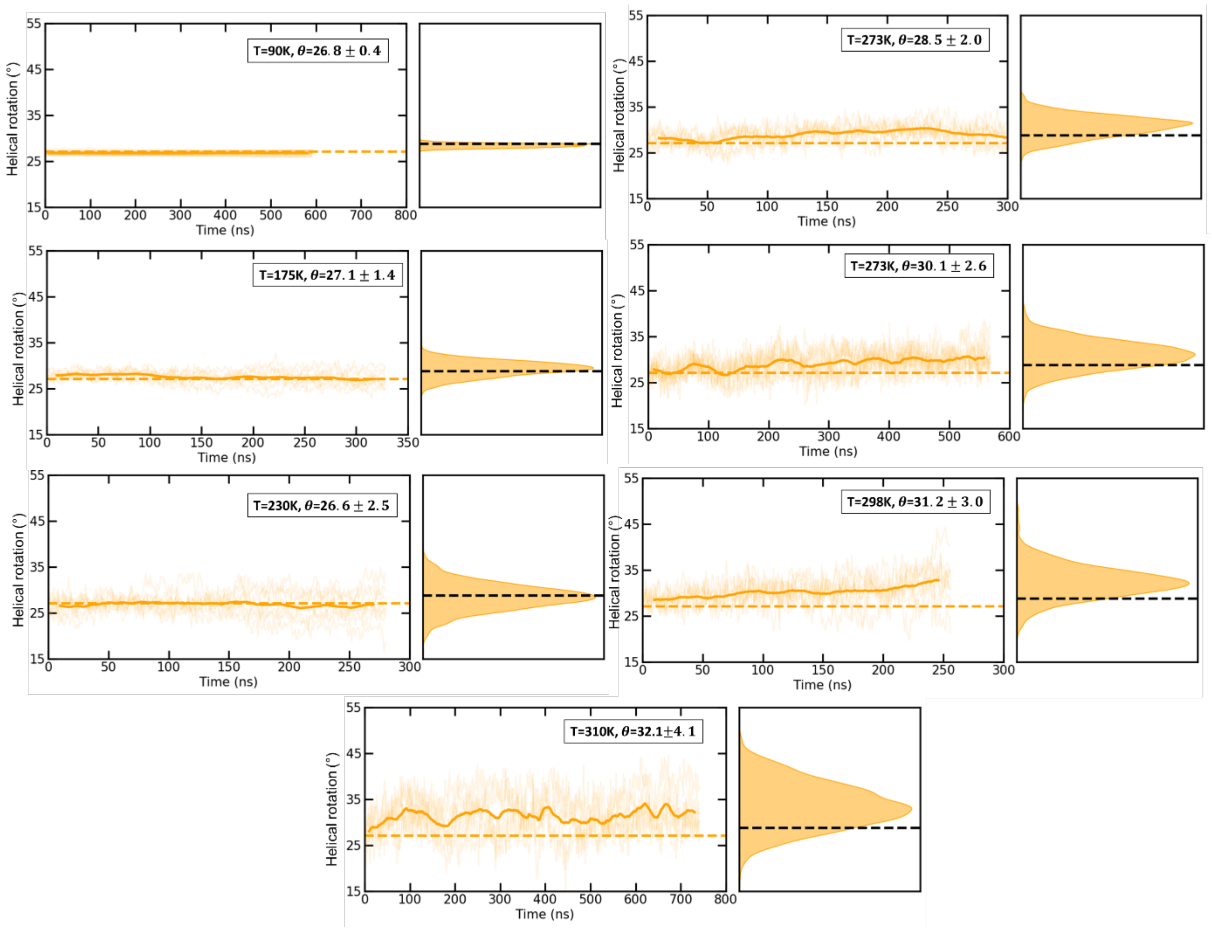


**Figure S4:** Time evolution of average dihedral angles and probability distribution of helical rotation angles of 8 internal subunits of 13-mer ADP-actin filaments at different temperatures. Text inserts are the averages and SDs calculated during the last 100 ns of the equilibrated trajectory.

**Section 4: Correlation between helical rotation angles, dihedral angles and lateral and longitudinal contacts.**

To understand the mechanism that allows for subunit twisting and fluctuations in the helical angle rotation, we measured the correlation between lateral and longitudinal contacts and the dihedral and helical rotation angles. The lateral and longitudinal contacts were measured within a cut-off of 3.5 Å. To calculate the correlation between lateral/longitudinal contacts and dihedral angles, we picked internal subunits 13 and 18 that consistently sampled dihedral angles greater than (or lesser) than -9° in the last 50 ns of the 1 microsecond trajectory. Since the dihedral angle and helical rotation angles were not correlated ($r^2 = 0.11$,Fig. S4A), we picked different internal subunits to calculate correlation between the helical rotation angles and lateral/longitudinal contacts. These subunits 10 and 15 had helical rotation angles consistently greater than (or lesser) than -162° continuously in the last 50 ns of the 1 μs trajectory. Hence, the data in Fig. S2 is clustered in two groups. Only 55% of the variation in dihedral angles can be attributed to loss of lateral contacts ($r^2 = 0.55$,Fig. S4B) and 60% of the variation in helical rotation angles can be attributed to loss of lateral contacts ($r^2 = 0.60$,Fig. S4C). The longitudinal D-loop interactions are more persistent than the lateral interactions, so the dihedral and helical rotation angles were not correlated with fluctuations in the lateral interactions (Fig. S4D,E). To summarize, (i) dihedral and short-pitch rotation angle are weakly correlated. i.e., subunits with more dihedral twisted conformation need more have more negative short-pitch rotation. (ii) dihedral angle and lateral contacts are weakly correlated. i.e., more twisted conformations occur when lateral contacts are lost. (iii) short-pitch rotation angle and lateral contacts are weakly correlated. i.e, subunits which less negative short-pitch rotation angle occur when lateral contacts are lost.

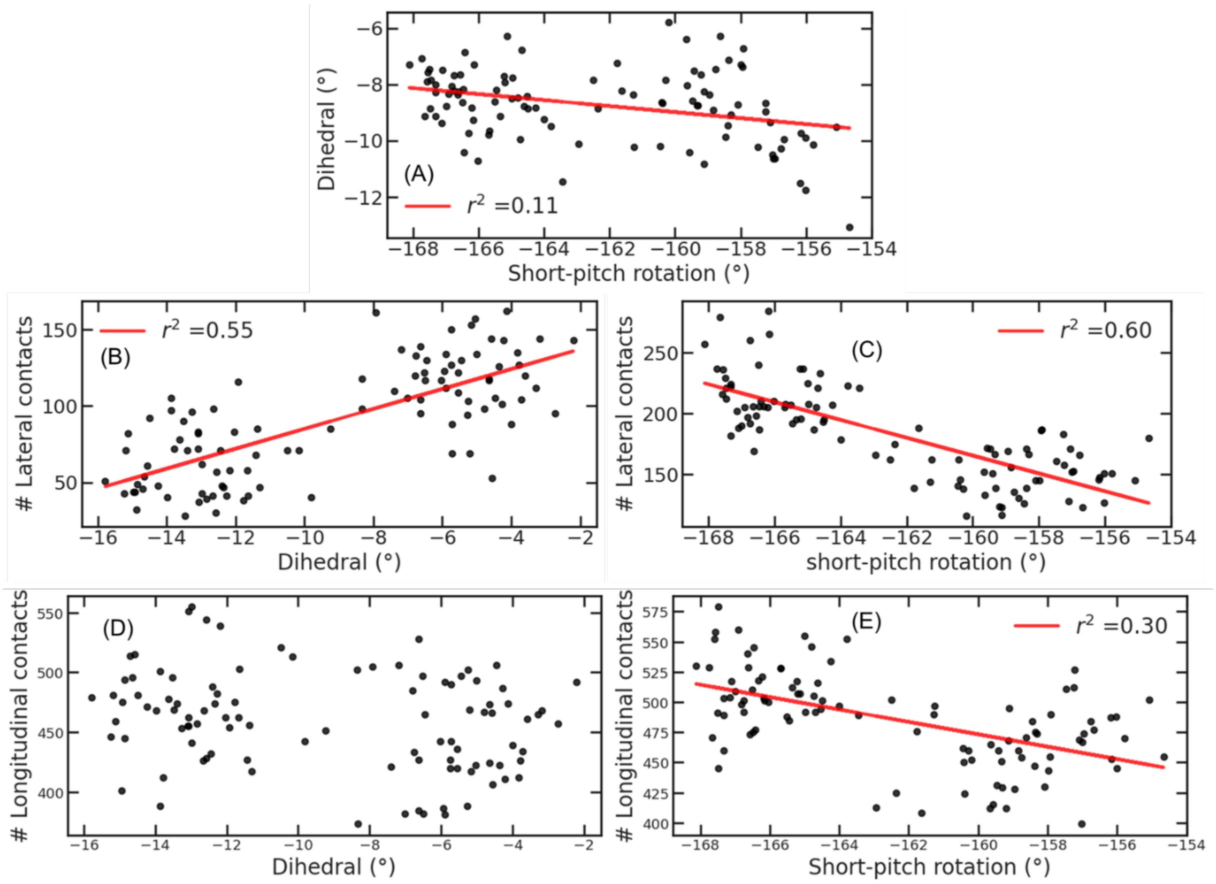


**Figure S5:** Correlation between helical rotation angles, dihedral angles and lateral and longitudinal inter-subunit contacts: (A) Correlation between dihedral angles and helical rotation angles sampled by the subunits. Correlation between lateral contacts with (B) dihedral angles and (C) helical rotation angles. Correlation between longitudinal D-loop contacts with (D) dihedral angles and (E) helical rotation angles.

### Section 5: Simulation of an ATP-actin filament with 13 subunits

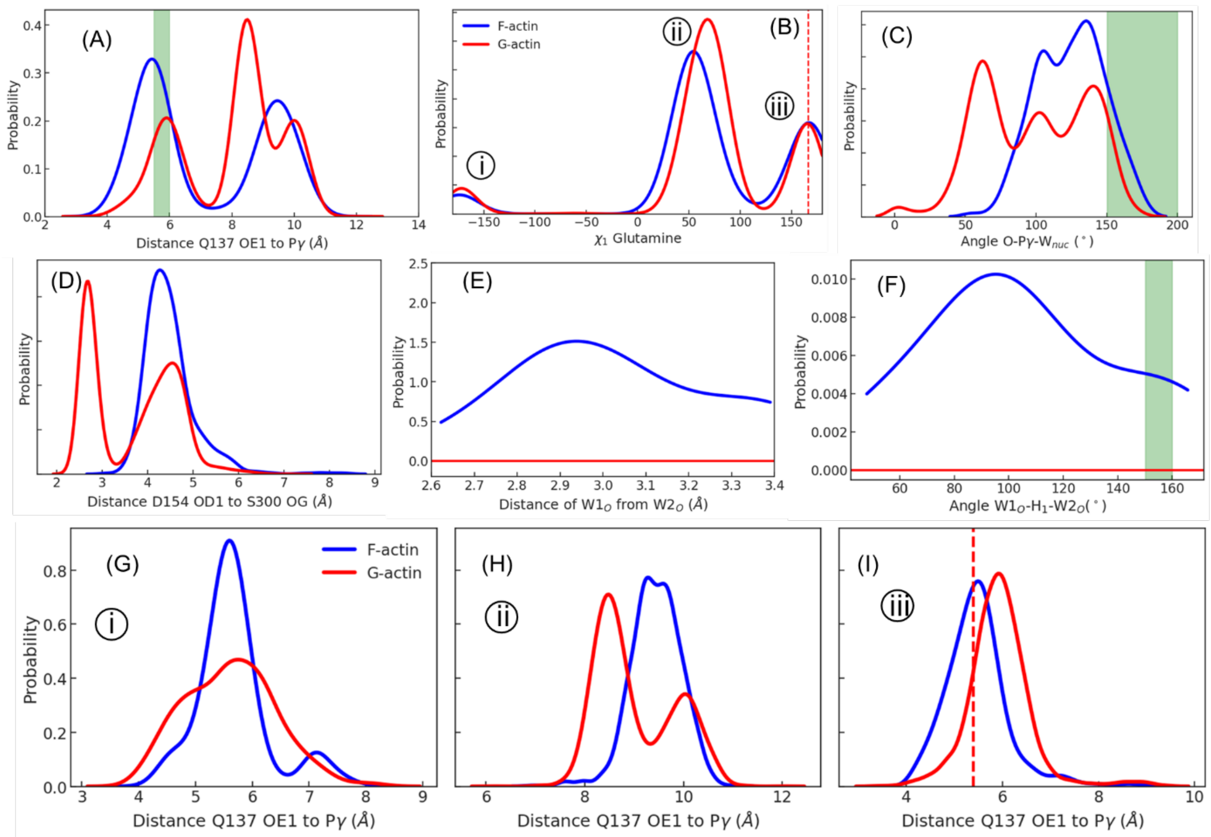


**Figure S6:** Replica 2 simulation of the ATP-actin filament with 13 subunits. (A) Probability distributions of distances between Q137 OE1 and γ phosphate of ATP in internal subunits of 13-mer ATP-actin filament during 200 ns and ATP-actin monomer during 4 µs of MD simulation. (B) Probability distribution of $\chi_1$ rotamers of Q137 in the internal subunits of an ATP-actin filament with 13 subunits sampled over the 200 ns trajectory and in an ATP-actin monomer sampled over 4 µs of MD simulation. Dashed line indicates the $\chi_1$ angle in cryo-EM reconstructed structure pdb id 8A2R. (C) The angle O-γ phosphate-W1 for subunits with distances of 5.5 Å to 6.0 Å between Q137 OE1 and γ phosphate. (D) Probability distribution of distance between OD1 of D157 and OD of S300. (E) Probability distribution of distance between W1 and W2. W2 is identified as a water located within 3.5 Å of W1 oxygen and coordinated to D157 and H161. (F) Probability distribution of the angle OW1-HW1-OW2. Angles that are optimal for hydrogen bonding are marked in green shaded region. (G-I) Probability distribution of distances corresponding to rotameric states (i)-(iii) of Q137 in panel B. Dashed line in (I) indicates the distance in cryo-EM reconstruction pdb id 8A2R.

In the second replica simulation: Condition 1 occurs during a fraction of 0.25 of the time in interior subunits of filaments (Fig. S6 A), condition 2 in only 0.16 of interior subunits satisfying condition 1 (Fig. S6 C), condition 3: in 0.4 of waters satisfying condition 1 and 2 and condition 4 for a fraction of 0.02 of the time in interior subunits of filaments. During this time, W2 has an optimal geometry to hydrogen bond with W1 for a fraction of 0.25 of the time in actin filament subunits (Fig. S6 E,F) . Combining these probabilities, the fraction of subunits in Mg-ATP-actin filaments with catalytically favorable waters in replica 2 is 0.25 x 0.16 x 0.4 x 0.02 x 0.25 = 0.00012. These rare subunits hydrolyze ATP at 0.3 $s^{-1}$/ 0.00008= 3750 $s^{-1}$.

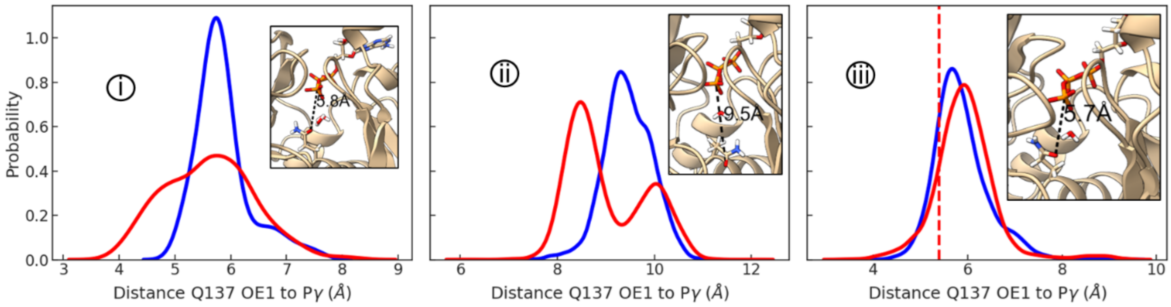


**Figure S7:** Probability distribution of distances corresponding to rotameric states (i)-(iii) of Q137 in Fig. 3D. Dashed line in (iii) indicates the distance in cryo-EM reconstruction pdb id 8A2R.

**Section 6: H161 in ADP-Pi filaments**

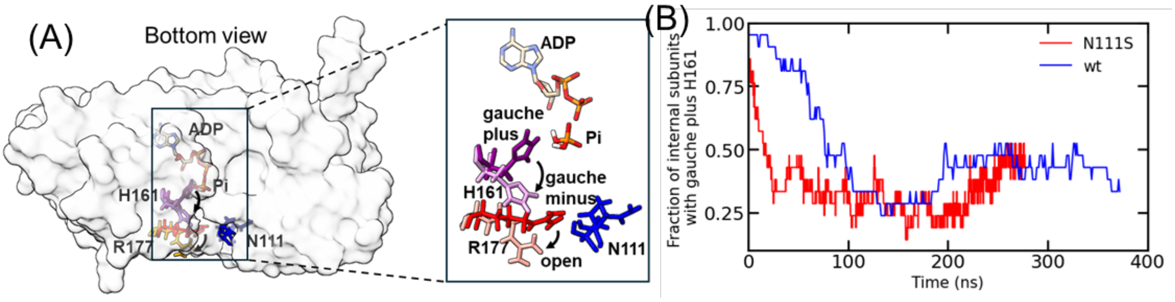


**Figure S8:** (A) Space filling model of bottom of a subunit in wild type-actin (light grey surface) with CPK representation of ADP-$P_i$, residue H161, N111 and residue R177. The two structures for each residue are the conformations at the beginning and after 200 ns of MD simulation. (B) Fraction of gauche plus rotameric conformations of H161 in internal subunits of wild type (blue) and mutant (N111S, red) filaments.

**Section 5: Comparison of actin subunits in the cryo-EM reconstitution and an MD simulation of a filament.**

As noted in (4) the twisted conformations of P and P-1 differ from the twisted conformation of actin monomers. Not only are the outer subdomains of subunit P rotated relative to the inner subdomains, but SD2 is also bent relative to SD1 (Fig. S3A).

Ribbon diagrams in Fig. S3 compare the conformations of the actin monomer, subunits in cryo-EM reconstructions and subunits 1 μs MD simulations of filaments with 27 subunits. Fig. S3A compares a crystal structure of an actin monomer aligned on the inner subdomains (SD3 and SD4) with subunits P, P-1, I and B in the cryo-EM reconstructions to illustrate how the orientations of subdomains SD2 and SD1 differ in monomers and filaments.

Figs. S3B-D compare the most extreme conformations of each actin subunit during MD simulations of the 27-subunit filament represented by the minimum and maximum dihedral angles in Fig. 1C. This variation is larger for the pointed subunit P than the barbed and the internal subunits.

In addition to reporting dihedral angels, we measured bending of SD2 relative to SD1 and SD4 relative to SD3 during 1 µs MD simulations of filaments with 27 subunits (Fig. S3E,F). We measured bending by aligning SD1 and SD3 at the first time point of the production run with subsequent structures. In subunit P, the large bend of SD2 relative to the cryo-EM structure resulted in an extreme twisted conformation (-23.2°), while bending of SD4 of subunit B resulted in mildly twisted conformation (-12.8°). SD2 bent less in internal subunits. The dashed lines in Fig. 2F show the orientations of SD1 and SD4 in the cryo-EM reconstructions.

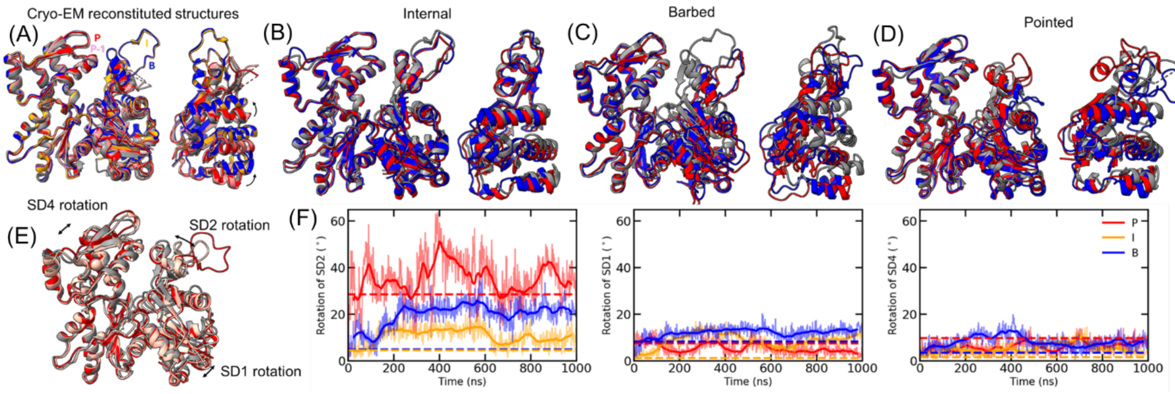


**Figure S9:** Comparisons of subunit conformations in different parts of actin filaments. (A) Comparison of ribbon diagrams of an actin monomer (grey, PDB ID: 3HBT) with filament subunits from cryo-EM structures aligned by subdomains 3 and 4. Filament subunit P is red (PDB ID: 8F8S), subunit P-1 is pink (PDB ID: 8F8S), an internal subunit is yellow (PDB ID: 8F8P) and subunit B is blue (PDB ID: 8F8R). Ribbon diagrams on the left show differences in SD1 and SD2. Ribbon diagrams on the right show the flattening by rotation of SD1 relative to SD3 (lower curved arrow) and bending of SD2 relative to SD1 (upper curved arrow). (B-D) Extreme conformations of the (B) internal, (C) barbed and (D) pointed subunits from the last 200 ns and the cryo-EM reconstructed structures. The extreme conformations and the cryo-EM reconstructed structure (grey) are aligned on SD3 and SD4 subunits. The red and blue ribbon diagrams are the most negative dihedral angle and largest dihedral angle for each of the subunit respectively. (E, F) Characterization of the rotation of SD1, SD2 and SD4 domains. (E) Ribbon representation of subunit P in the cryo-EM reconstitution (grey), first fame (red) and at the end of 1 µs MD simulation (pink) aligned on SD3 of the first fame of production (after equilibration) to show the rotation of SD1, SD2 and SD4 with respect to first frame of production. Arrows indicate rotation of each subdomain with respect to the reference first frame in red. (F) The three panels show the time evolution of rotation of SD2 (left), SD3 (middle) and SD4 (right) with respect to first frame of production in the 1 µs simulation. Dashed lines are the angles in the cryo-EM models aligned on SD3 of the first frame of production after pre-equilibration.

**Section 6: Effects of filament length on the behavior of the ends during MD simulations.**

To study the effects of filament length on the actin subunit conformations, we ran MD simulations of actin filaments with 7 or 13 subunits to compare with filaments of 27 subunits. The 7-mers correspond to half of the cross over distance, 13-mer to the half-pitch (36 nm) and 27-mer to one full turn of the filament double helix (72 nm).

**Pointed end simulations:** Figures S4-S6 present the analysis of pointed ends during three replica runs: Fig. S5 for the for the 7-mer, Fig. S6 for the 13-mer and Fig. S7 for the 27-mer. The graphs in all three figures show the time evolution of (upper row) dihedral angles of subunit P as defined in Fig. 1C, (middle row) distances between the atoms that form a salt bridge between NE of R62 in P-1 and OE1 or OE2 of E270 in P.  and (bottom row) number of contacts within 3.5 Å between P and P-1 and I and I-1 subunits. The vertical orange lines mark the time point when salt bridge interactions begin to form between R62 to E270. The horizontal purple line notes a distance of 5 Å, which is favorable for salt bond formation. The ribbon diagrams in all three figures show the pointed ends at the ends of the simulations. In all 3 replica runs the P-1 subunit and its D-loop bends towards the P subunit, limiting interactions of the D-loop with the barbed end of an incoming actin monomer.

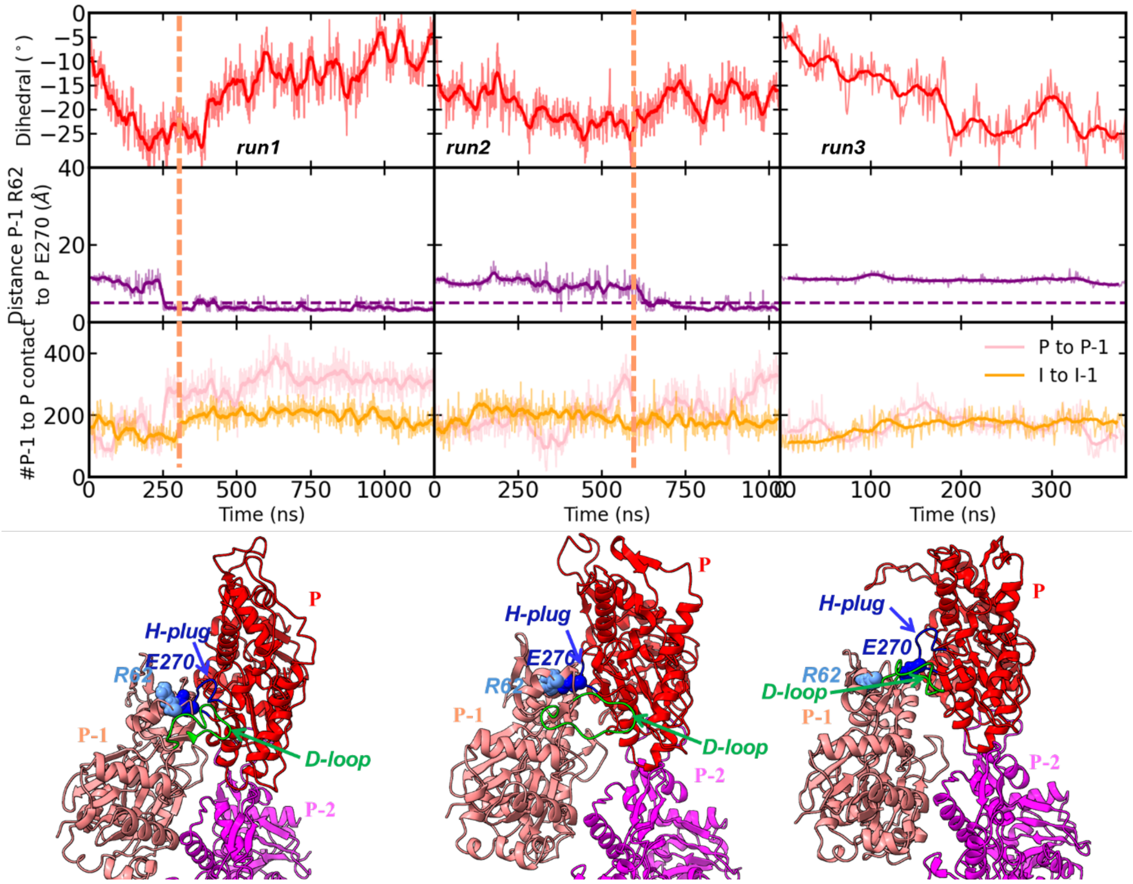


**Figure S10:** Pointed end from ~2.5 μs of MD simulations of a 7 subunit filament.

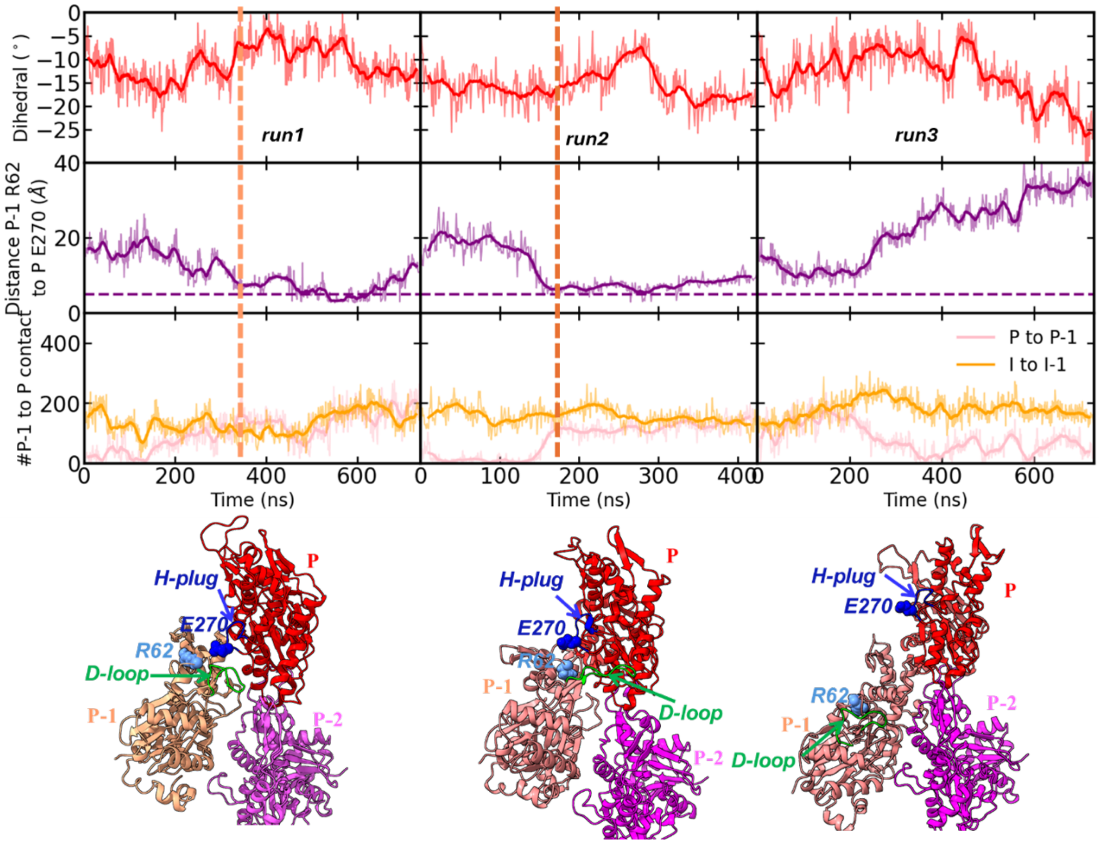


**Figure S11:** Pointed end from ~1.9 μs of MD simulations of the 13 subunit filament. In all three replica runs the P-1 subunit bent towards P subunit for much of the simulation, limiting binding to the barbed end of incoming actin monomer and dissociation of subunit P. In the 3rd replica the contacts between P-1 and P subunits were lost after about 400 ns, which may favor dissociation of subunit P.

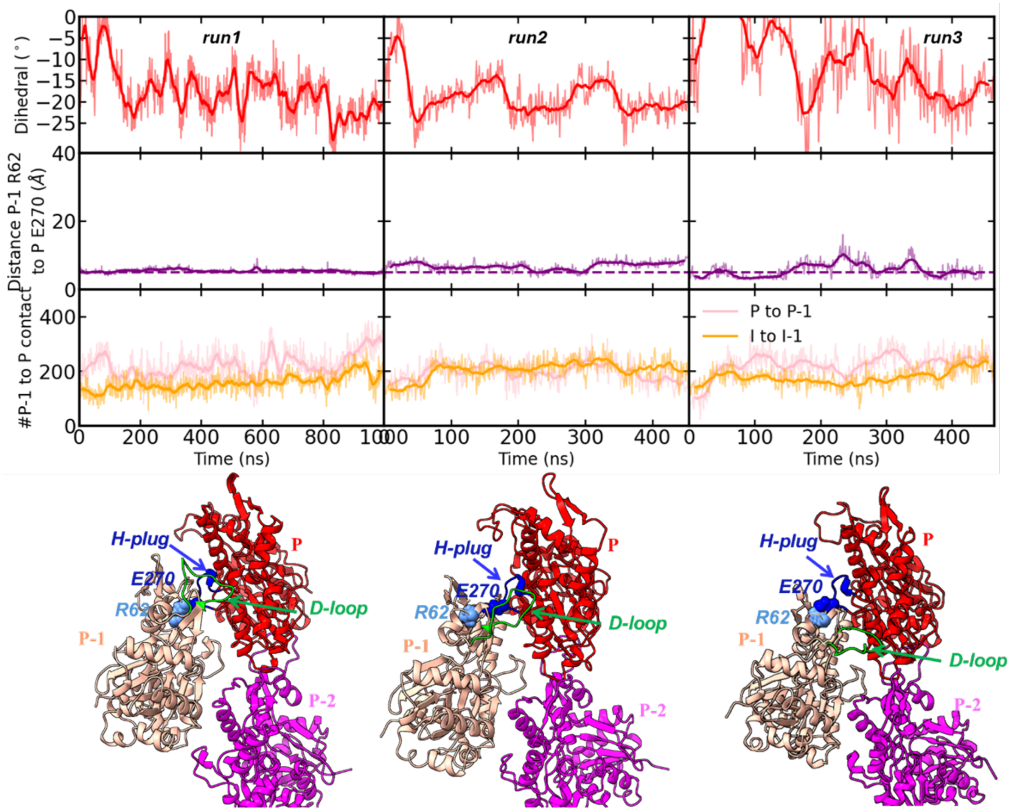


**Figure S12:** Pointed end from ~1.9 μs of MD simulations of a 27-mer subunit.

**Barbed end simulations:** Fig. S8 compares the dynamics of barbed ends with internal subunits during replica MD simulations of filaments with 7, 13 or 27 subunits.

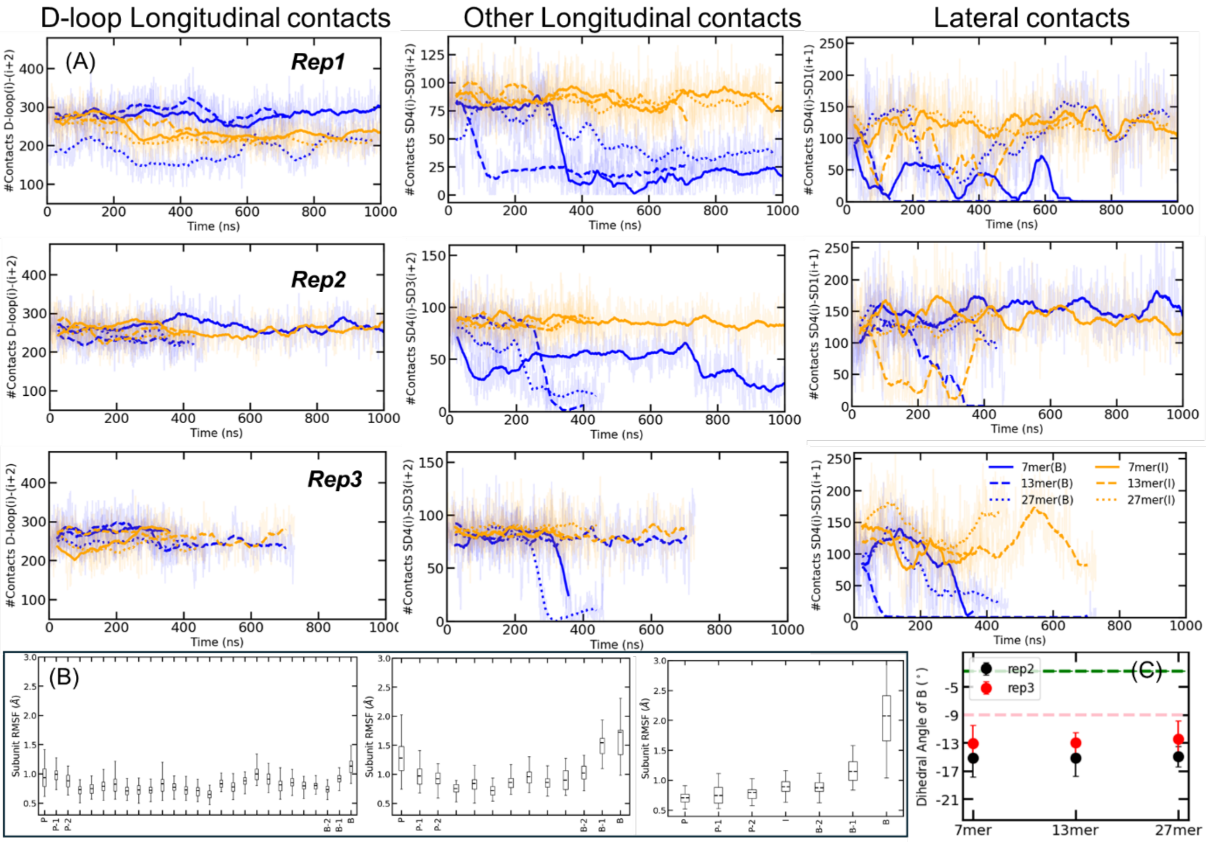


**Figure S13:** Dynamics of barbed end and internal subunits during replica simulations 2 and 3 of filaments with 7, 13 or 27 subunits. (A) Time courses of the number of longitudinal and lateral contacts within a cut-off of 3.5 Å for internal subunits (orange) and barbed end subunit B (blue). Longitudinal contacts were calculated between and the D-loop of (i) with barbed end groove of (i+2) and other contacts between SD4(i) and SD3(i+2). Lateral contacts are calculated between SD4 (i) and SD1 (i+1). (B) Root mean square fluctuations of the centers of mass of subunits in (left) 27-mer, (center) 13-mer and (right) 7-mer filaments during the last 200 ns of the simulations. Edges of the box plot represents first and third quartiles and the horizontal line within each box are the mean RMSF values. Whiskers extend to minimum and maximum RMSF sampled. (C) Average and standard deviation of dihedral angles of the barbed subunit of actin filaments with 7, 13 and 27 subunits measured in the last 200 ns of second and third replica runs of the simulations. Dashed green line at -3.0° corresponds to the dihedral angle of subunit B in the cryo-EM reconstruction (pdb id: 8F8R). Dashed pink line at -9.0° corresponds to the average dihedral angle of internal subunits in 310 K MD simulation.

Throughout all three simulations of filaments with 7, 13 and 27 subunits the internal subunits maintained their longitudinal contacts (Fig. S8A yellow lines in columns 1 and 2). On the other hand, the lateral contacts were less stable (Fig. S8A yellow lines in column 3).

The D-loop of subunit B also maintained its contacts with subunit B-2 (Fig. S8A, blue lines in the first column), while SD4 of subunit B lost longitudinal contacts with SD3 of subunit B-2 (Fig. S8, blue lines in the second column) and lateral contacts with subunit B-1 (Fig. S8, blue lines in the third column) stochastically for variable durations (Fig. S8A). Without these variable contacts subunit B was more flexible and adopted slightly more twisted conformations compared to the average dihedral angle of internal subunits -9.0° (Fig. S8C).

Table S3 summarises the dihedral angle subunit B for the replica runs of 7, 13 and 27 subunit filaments when the lateral contacts between SD4(i) and SD1(i+1) are intact/lost.

The lateral contacts are stable for longer in 27 subunit than 13 subunit and 7 subunit filaments (Fig. S8A). Lateral contacts are lost 20% of the time in the total 1.9 μs of simulation of the 27 subunit filament, 75% of the time in the total 1.89 μs of simulation of the 13 subunit filament and 25% of the time in the total 2.59 μs of simulation of the 7 subunit filament This is reflected in the RMSD fluctuations of subunit B in filaments with 7 and 13 subunits vs. 27 subunits (Fig S8B). In 7 and 13-mer filaments, the RMSF of center of mass of subunit B is greater than 1.5 Å, while for the 27-mer the RMSF is less than 1.2 Å. B subunit RMSF scaled inversely with filament length.

**Table S3:** Summary of dihedral angles of subunit B for filaments with 27, 13 and 7 subunits in 3 replica simulations. Dihedral angle (SDs) were calculated during the time segments when lateral contacts were intact or lost.

| Filament | rep | Intact lateral contacts | | Dissociated lateral contacts | |
|---|---|---|---|---|---|
| | | Time (ns) | Dihedral (SD) | Time (ns) | Dihedral |
| 27-mer | 1 | 600 – 800 | -11.4 (3.3) | 250-400 | -16.1 (2.5) |
| | 2 | 0-200 | -10.6 (3.9) | | |
| | 3 | 100-200 | -8 (1.5) | 350-450 | -13.0 (3.5) |
| 13-mer | 1 | | | 400-600 | -18.0 (2.5) |
| | 2 | 0-150 | -11.0(3.8) | 400-600 | -15.7 (2.7) |
| | 3 | | | 300-400 | -14.1 (2.5) |
| 7-mer | 1 | | | 800-1000 | -21.2 (2.2) |
| | 2 | 300-500 | -11.1(2.3) | | |
| | 3 | 100-200 | -10.1 (1.9) | 350-400 | -13.4 (3.1) |

**Section 7: Convergence of free energy sampling.**

To access the convergence of free energy during well-tempered metadynamics simulations for the terminal barbed end, internal, and terminal pointed end subunits, we monitored the stabilization of the potential of mean force (PMF, aka free energy surface), finding that further sampling does not significantly change the calculated PMF. The PMF was obtained via a c(t) reweighting procedure (5). The simulations were reweighted on the dihedral angle collective variable used for biasing. The PMF was calculated at the first and second instance when the whole range of phase space is sampled. The time this occurred was 500 ns for the barbed and pointed end subunits and 250 ns for the internal subunits (vertical red bashed lines in the top row of Fig. S14). The PMF obtained in the two instances was compared by aligning their minimum to zero (bottom row Fig. S14).

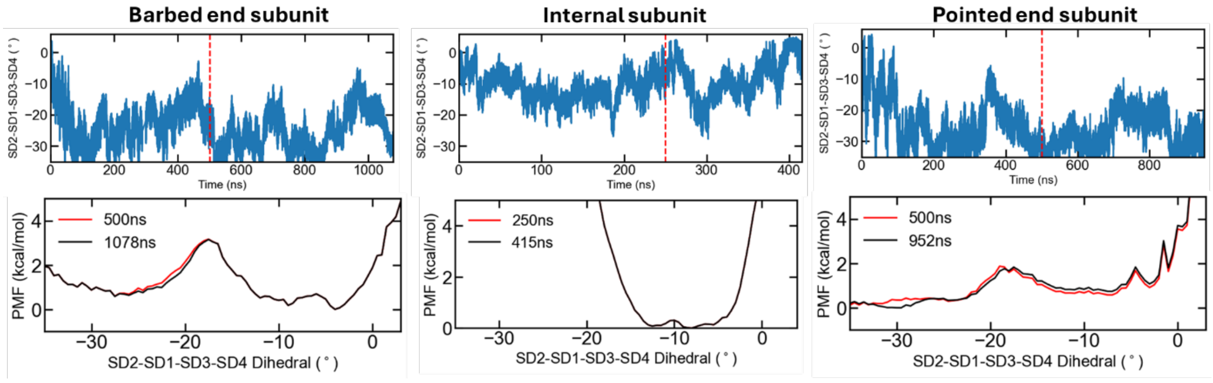


**Figure S14:** Convergence of free energy sampling for (left) terminal barbed end subunit B, (center) internal subunits and (right) terminal pointed end subunit P for 7-mer actin filaments. Top row shows the dihedral angles sampled during the well-tempered metadynamics simulation and the bottom row shows the reweighted Free Energy Surface (FES) projected on the dihedral angle with different time points. The dashed red line in the top row indicates the time at which the whole range of dihedral values were sampled for the first instance. These time points are used to compare the reweighted Potential of Mean Forces (PMFs) shown in bottom row.
